# Gram-negative outer membrane proteins with multiple β-barrel domains

**DOI:** 10.1101/2021.02.17.431615

**Authors:** Ron Solan, Joana Pereira, Andrei N. Lupas, Rachel Kolodny, Nir Ben-Tal

## Abstract

Outer membrane beta barrels (OMBBs) are found in the outer membrane of Gram-negative bacteria and eukaryotic organelles. OMBBs fold as antiparallel β-sheets that close onto themselves, forming pores that traverse the membrane. Currently known structures include only one barrel, of 8-36 strands, per chain. The lack of multi-OMBB chains is surprising, as most OMBBs form oligomers and some function only in this state. Using a combination of sensitive sequence-comparison methods and co-evolutionary analysis tools, we identify many proteins combining multiple beta barrels within a single chain; combinations that include 8-stranded barrels prevail. These multi-barrels seem to be the result of independent, lineage-specific fusion and amplification events. The absence of multi-barrels that are universally conserved in bacteria with an outer membrane, coupled with their frequent *de novo* genesis suggests that their functions are not essential, but rather beneficial in specific environments. Adjacent barrels of complementary function within the same chain may allow for new functions beyond those of the individual barrels.

## Introduction

Outer membrane beta barrels (OMBBs) are an important class of membrane proteins in Gram-negative bacteria, mitochondria, and chloroplasts [2, 3]. In bacteria, OMBBs are the most common family of outer membrane proteins and their functions are very diverse (e.g., adhesion, pilus formation, specific and nonspecific forms of import and efflux, proteolysis, even outer membrane protein assembly). Structurally, OMBBs are closed β-sheets of antiparallel β-strands, forming pores that traverse the membrane. In the majority of the cases, this sheet is formed from a single protein chain, but in some cases it can result from the assembly of smaller β-sheets, as in trimeric autotransporters [4]. With the exception of the mitochondrial 19-stranded porins [5], OMBBs have an even number of strands, and OMBBs with between 8 and 36 strands have been described so far [6, 7]. For their biological activity, many OMBBs form complexes with each other, as for example, the outer membrane phospholipase A (OMPLA) homodimer [8, 9], the trimeric porins [10, 11], and the type-9 translocon heterodimer [7].

Previous studies highlighted the common evolutionary origins of OMBB proteins [6, 12, 13]. Remmert et al. [12] argued that all OMBBs in Gram-negative bacteria evolved from a single ancestral subunit of two β-strands, arranged as a hairpin. Their study, and a later work by Franklin et al. [6], showed that the diverse structures of OMBBs evolved through amplification and recombination of these hairpins, and the accretion of mutations. These analyses point to the important contributions of duplication and fusion events to the evolution of OMBBs. In an experimental study, Arnold et al. [14] showed that the fusion of two β-barrels connected by a short linker can yield a single barrel of twice the size, demonstrating that concatenating hairpins may result in single, larger barrels. Given the importance of amplification for the genesis of new β-barrels, it is notable that, in spite of the propensity of OMBBs to form oligomers, and the fact that some OMBBs function solely in complex with other OMBBs, naturally-occurring proteins with multiple barrel domains (referred to herein as “multi-barrel” proteins) have not yet been identified (though the possibility of their existence has been acknowledged, e.g., by Reddy and Saier[13]).

We sought to expand the current repertoire of known OMBB architectures and provide insights to their evolution by searching for proteins with multiple OMBB domains. To identify novel protein architectures, one must search beyond the PDB [15], in the structurally uncharacterized space curated in sequence databases [16]. Using a combination of state-of-the-art sensitive sequence-comparison methods and co-evolutionary analysis tools, we searched the UniRef100 and the NCBI non-redundant (nr_bac) databases for sequences with more than one non-overlapping match to any OMBB family of known structure. After classification and annotation of the identified sequences, we were able to predict many putatively new OMBB families with novel strand-topologies, but most importantly, with a wide variety of multi-barrel domain combinations. Based on the function of their closest single-barrel homologs, we discuss putative biological roles for some of the multi-barrel families. Our findings highlight that OMBBs have a richer repertoire of architectures than previously known, expanding our current knowledge of these proteins, providing hints about their evolution, and revealing more than 30 new architectures of Gram-negative surface proteins.

## Results and Discussion

To identify putative multi-barrel proteins and single barrels with yet unknown architectures, we followed two parallel and complementary approaches. In one approach, we used HMMER [17] to align Hidden Markov models (HMMs) of all single-barrel proteins of known structure to the sequences in UniRef100 [16] and searched for cases with multiple matches along their sequence. In a second approach, we used PsiBlast [18] to search the bacterial protein sequences in the non-redundant sequence database at NCBI (*nr_bac*), starting from representative sequences of all OMBB families of known structure. Both approaches resulted in a similar set of proteins, which were combined, classified, and annotated using state-of-the-art sequence analysis tools.

We grouped the protein sequences that we found based on their global sequence similarity and denote these multi-barrel families (MB-families). In some cases, we further grouped MB-families that share some sequence similarity, but not necessarily over their entire chains, and denoted these multi-barrel superfamilies (MB-superfamilies). An alternative grouping of the MB-families is based on their predicted structure, denoted as multi-barrel architecture (MB-architecture). The MB-architecture annotates the sizes of the barrel domains, ordered from the N-terminal domain to the C-terminal domain. For example, the MB-architecture of a protein composed of a 16-stranded N-terminal barrel and a 12-stranded C-terminal barrel is 16-12. The search and annotation procedures are described in detail in the Methods section. The seeds and MB- architectures found are available online at http://trachel-srv.cs.haifa.ac.il/rachel/MOMBB/.

### Large variety of multi-barrel architectures

We identified 12,643 unique proteins with a new architecture and provide an overview of this large set using CLANS’s coarse clustering based on local sequence similarity (Figure S1). A finer clustering, based on global similarity and manual inspection, allowed separating these coarse clusters into 186 MB-families, each with a single MB-architecture. Of these MB-families, available at http://trachel-srv.cs.haifa.ac.il/rachel/MOMBB/, 79 have four or more proteins (Table S1). Size distribution shows many small MB-families and a few large MB-families of ~1000 proteins or more (Figure S2).

Annotation of representative proteins from each cluster suggests that 34 novel MB-architectures are represented in our set (Figures 1 and S3). These mostly comprise proteins with multiple non-overlapping, full-length matches to known OMBBs, indicating concatenation of known OMBBs. The linkers connecting these matches in representative sequences from each MB-family (the ones used to annotate the architectures in Figure 1) provide a clear delimitation between the putative barrels, with a median length of 22±13 residues (Figure S4). These findings suggest that a large variety of multi-barrel domain architectures exist in nature, far beyond the previously documented repertoire of OMBBs.

**Figure 1:**
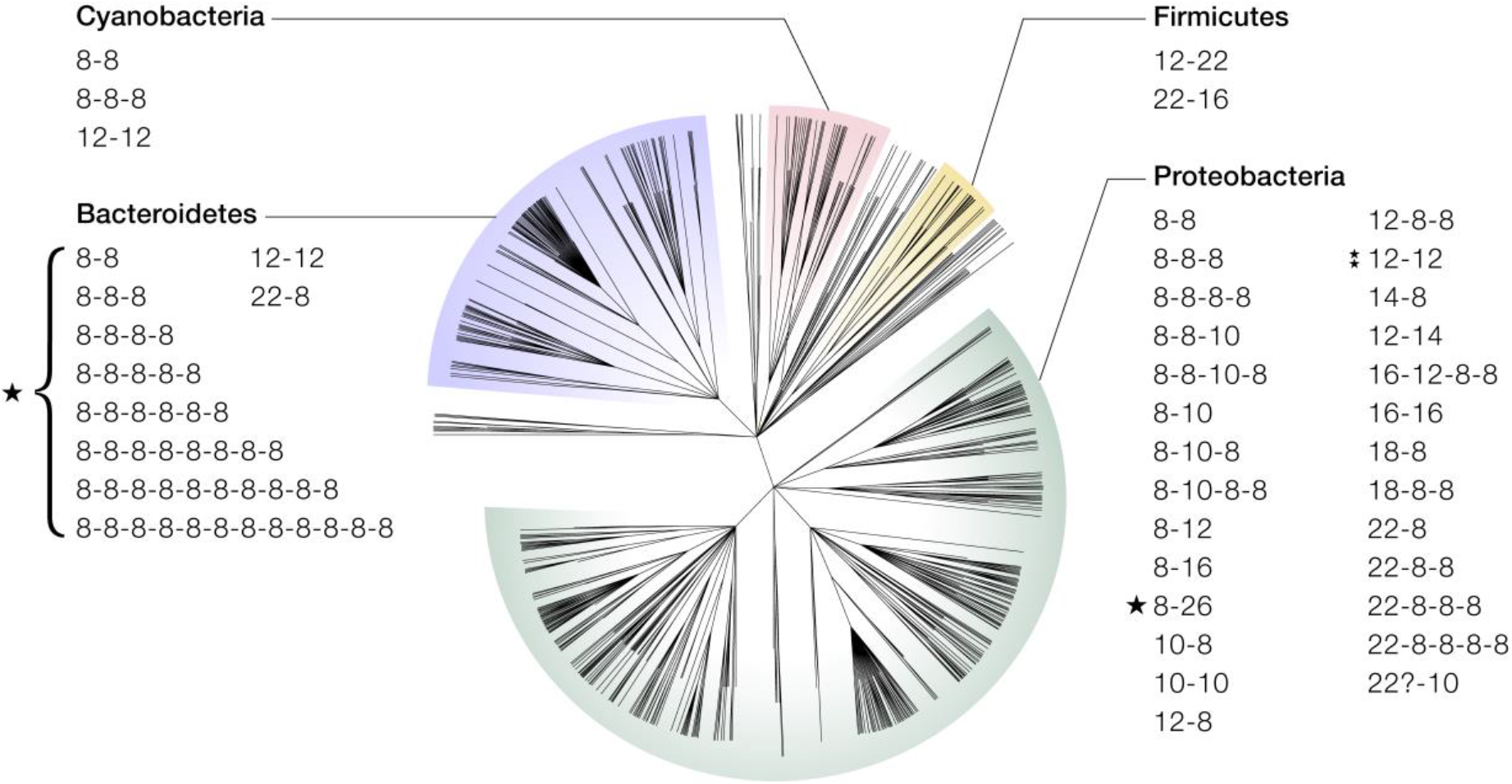
Taxonomic classification of the multi-barrel architectures. All the species with MB architectures were collected, and their taxonomic tree was composed from the NCBI taxonomy database. The MB-architectures that are manifested in each of the major clades are listed, with asterisks marking the ones that are discussed in the main text. The tree was rendered using Dendroscope[1].

Most of the architectures involve an 8-stranded barrel, i.e., are either two (or more) concatenated 8-stranded barrels, or an 8-stranded barrel concatenated with a barrel of different size (Figures 1 and S3). All but one of the architectures with at least three barrels are composed of multiple 8-stranded OMBBs (Figures 1 and S3), or a combination of these and exactly one barrel of another size (of 10, 12, 16, or 22 strands). Some architectures are two barrels with the same number of strands: these can be either a repeat, as in the PLA1-PLA1 (12-12) architecture, or two barrels from different OMBB families, as in the FhaC-Porin (16-16) architecture. Not all known single-barrel topologies appear in multi-barrels: we did not find any multi-barrel architecture with a 24-stranded barrel or with a 36-stranded barrel.

Grouping by the taxonomy of their respective bacteria shows that the new multi-barrel architectures are present mostly in Proteobacteria and Bacteroidetes, although a few are in Cyanobacteria, Firmicutes, Fusobacteria, and other phyla (Figure 1).

We inspected the distribution of species within each MB-family. Some MB-families include only a few homologous proteins in a single species. All MB-families save one, are endemic to a single phylum (with up to only 2% proteins in other phyla). Figure 2 shows an example of two different distributions of two clusters: the 18-8-8 MB-family (marked with red circles) has 48 proteins but is spread among more Proteobacterial clades than the 22-8 MB-family (marked with yellow circles), which has 99 proteins.

**Figure 2:**
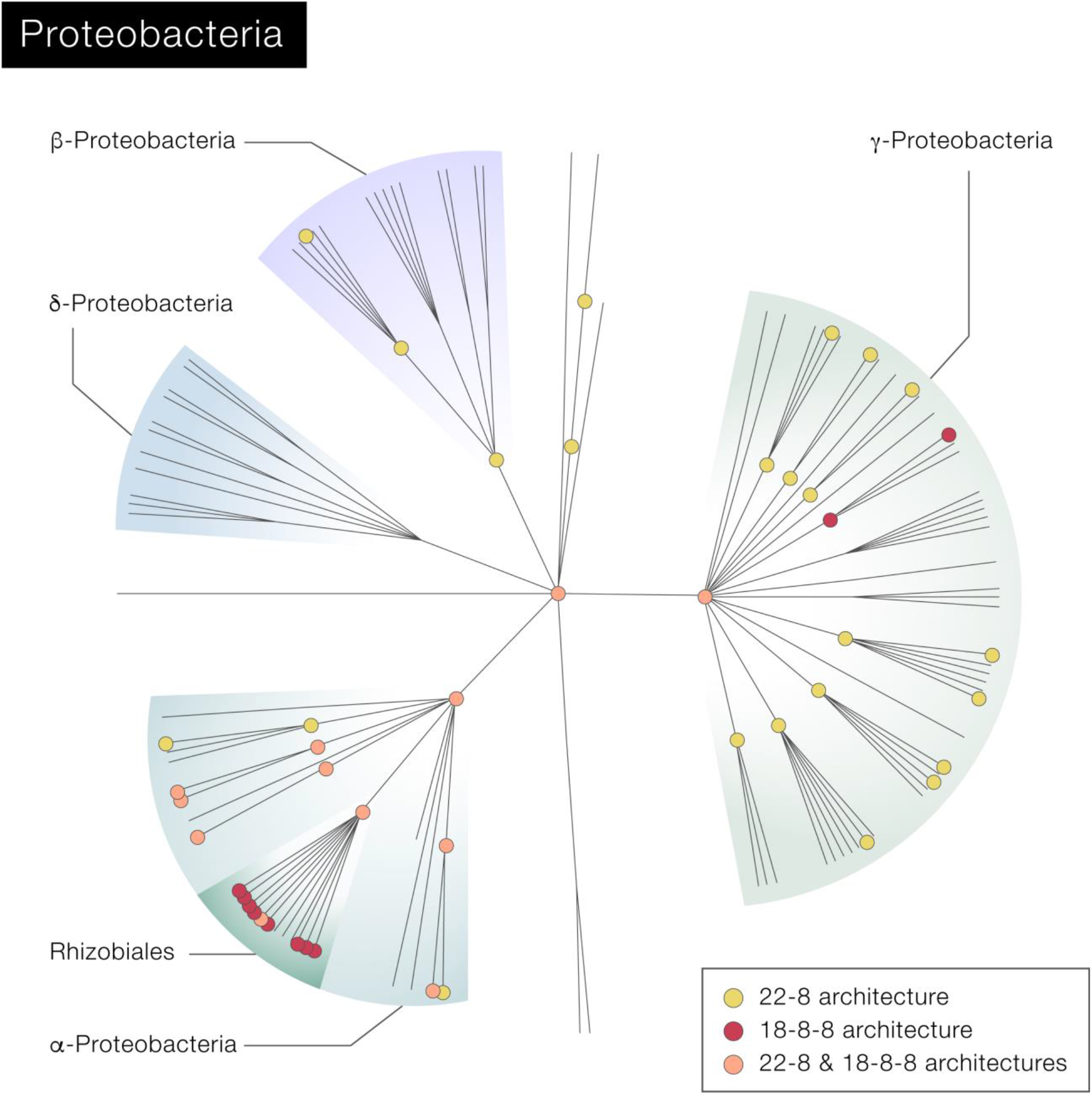
The taxonomic tree of the MB-families in Proteobacteria with multi-barrel proteins. Clades with an architecture of 22-8 are marked with a yellow circle, clades with an architecture of 18-8-8 are marked with a red circle, and clades with both architectures are marked with an orange circle. Even though the 22-8 MB-family (marked in yellow and orange) has twice as many representatives as the 18-8-8 MB-family (marked in red and orange), it appears mostly in Rhizobiales, which are α-Proteobacteria, while the proteins of the 18-8-8 MB-family are present in many orders in α, β, and γ Proteobacteria. The tree was rendered using Dendroscope [1].

### Additional architectures

Other instances, beyond those that clearly include multi-barrels, show only partial and/or overlapping matches to known OMBBs, representing putative new MB-families. Within these, we identified candidates for architectures with 10, 12, 16, and 34 strands, as well as an MB-superfamily of large OMBBs with at least 40 strands, larger than any OMBB reported to date. Some multi-barrels in our set include other domain families that are not integral to the membrane, some well characterized (such as POTRA domains) and others without any clear homology to a known family (Figure S1).

### Contact map predictions support the multi-barrel nature of the protein chains

The sequence matches for the architectures described above are generally full-length matches to known OMBBs, separated by extended linkers. Nevertheless, the connected barrel domains might still fold into larger fused barrels [14]. We use contact map predictions to examine this.

When there are many homologous and variable proteins sequences that fold into the same structure, computational tools can predict the two-dimensional residue-residue contact map of their structure [19, 20]. Because the prediction of the contacts exploits the correlated mutation signals in the multiple sequence alignment, a homologous protein with a known structure that will serve as a template is not required. In theory, correlations between mutations are a consequence of compensatory changes between residues in close physical contact [21]. In practice, however, correlated mutations are also observed between residues that are not in contact. To extract only true contacts from correlated mutations, state-of-the-art contact map predictors employ deep networks [22]. Herein, we generated contact maps using the RaptorX software tool, which the 2016 CASP12 and 2018 CASP13 experiments identified as an accurate predictor [22, 23].

Accurate prediction of contact maps requires multiple sequence alignments (MSA) of many diverse and homologous sequences [5, 19]; when only a few homologs are available, predictions tend to be of low quality. For example, 148 effective homologs were needed to reach an accuracy of 0.55 (in a 0-through-1 scale) in the top L/5 medium-range contacts in membrane proteins (L being sequence length) [19]. Only 13 MB-families have more than 148 homologous proteins, and even a smaller number have over 148 *effective* homologs (i.e., after redundancy cleaning). Nevertheless, we predicted contact maps for all MB-families with at least 50 proteins, to find cases where despite the small number of homologs, the signal is strong enough for accurate predictions. The anticipation was that in most of them there would be no signal.

To filter the many expected contact maps with no strong signal, we defined strict criteria for contact maps that we consider accurate. In proteins with one or more barrel domains, we expected that (e.g., Figure 3): within each of the barrels, there would be contact signals near the diagonal between every two consecutive strands, allowing to infer the number of strands in the barrel even if the predictions are noisy; there would be a barrel-closing signal far from the diagonal, indicative of contact between the N− and C-terminal strands, which are adjacent in the folded barrel; and (3) there would be no significant contacts between strands in different barrels, or contacts between non-adjacent strands in the same barrel. With these criteria in mind, we tried to predict contact maps with RaptorX for the 23 multi-barrel MB-families containing more than 50 protein sequences.

**Figure 3:**
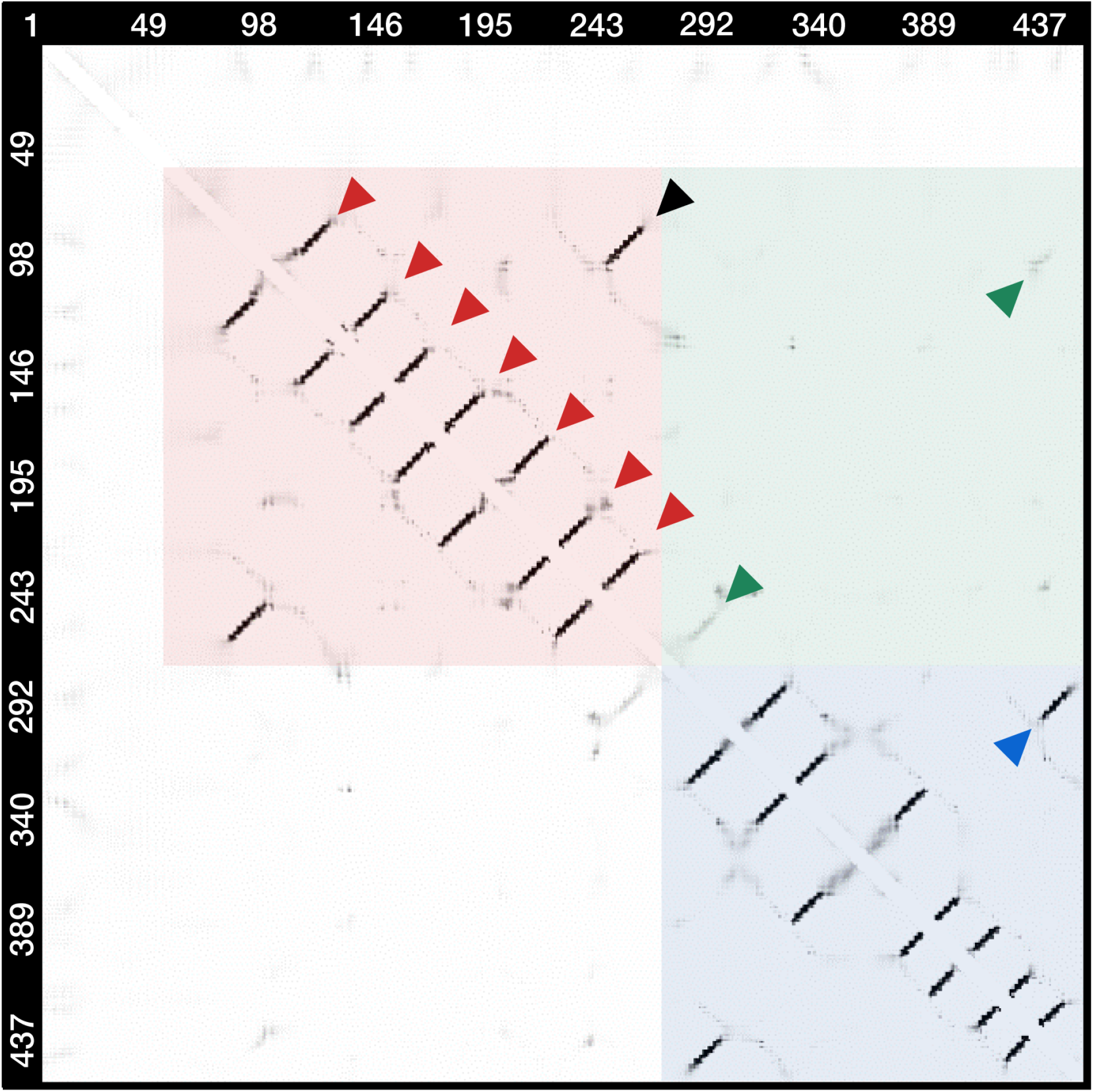
RaptorX prediction of the contact map of MB-family 001, comprising 1,959 homologous proteins, manifesting an 8-8 architecture (a pair of 8-stranded barrels). This predicted MB-architecture is strongly supported by the clear (dark) signals for contacts between the spatially adjacent beta-strands of the N− and C-terminal barrels within the red and blue frames, respectively. That there is, in essence, no signal within the green frame is a clear indication that the alternative 16-stranded barrel architecture is much less likely.

Figure 3 shows an example of a contact map prediction, supporting the hypothesis that the 1,959 homologs of MB-family 001 have two separate 8-stranded barrel domains. Contact maps are, by definition, symmetric; thus, we focus our discussion on the triangle above and to the right of the diagonal. Looking at the first part of the predicted contact map (~200 residues; red square) we see seven lines perpendicular to the main diagonal (red arrows). Each of these diagonal lines describes contacts between two consecutive strands, with residues of increasing number in the first strand in contact with residues of decreasing number in the second strand. Thus, the seven lines describe contacts between strands 1-2, 2-3, 3-4, …, 7-8. Finally, contacts between strands 8 and 1, in the upper right corner and marked by a black arrow, manifest a barrel-closing signal. The second barrel domain (enclosed in a blue square) has a similar pattern, with contacts between strands 8 and 1, i.e., barrel-closing signal, marked by a blue arrow. Beyond these clear patterns, there are only a few weak correlations in the predicted contact map, which we consider to be noise. In particular, the predicted signals of contacts between the two barrels (marked by the green arrows) are very weak. The proteins in this MB-family have two segments homologous to 8-stranded barrels. Theoretically, rather than forming two separate barrels, they could fuse to form a single large barrel. Since there are sequence matches to all the strands, the most likely single barrel would comprise 16-strands. In this case, we would expect to see a contact between strands 8 and 9, and between strands 1 and 16 (marked by green arrows), which are only weakly observed in the predicted contact map. Namely, the contacts supporting the 8-8 architecture hypothesis are much stronger than the contacts supporting the 16-strand hypothesis, and by visual inspection we conclude that the contact map supports the prediction of two consecutive 8-stranded barrel domains. Although RaptorX outputs contact probabilities, the dependencies among them prevent us from combining the contact probabilities for the strands in a mathematically meaningful way. Despite that, just as the example demonstrated in Figure 3, the visual signal is sometimes clear enough to identify the MB-architectures.

Figures S5 and S6 show the predicted contact maps of two more MB-families that clearly support their predicted MB-architectures: MB-family 007 with 8-8-8 architecture (Figure S5; although a barrel closing signal for the C-terminal barrel in missing), and MB-family 015 with 22-8 architecture (Figure S6). Similarly, Figure S7 shows the predicted contact map of MB-family 002, supporting a predicted single barrel, larger than previously observed. Indeed, the RaptorX contact map confirms the presence of a barrel with at least 40 beta strands. Altogether, predicted contact maps support four previously undocumented architectures (Figures 3, S5, S6 and S7). However, contact maps do not always yield clearly interpretable signals, even when there are many homologues. For example, in MB-family 003, predicted to form a 8-8 architecture, there are 813 sequences, but the contact map does not unequivocally support a single architecture (Figure S8). Another case is MB-family 000, with 4,187 homologues, predicted to form a 12-12 architecture. The corresponding contact map clearly supports the N- but not the C-terminal barrel domain (Figure S9). This could indicate that the correlated mutation signal for the C-terminal barrel is particularly weak, or that it is not a barrel domain.

Overall, the predicted contact maps provided further support for four novel barrel architectures. Three of these are multi-barrel architecture, 8-8 (Figure 3), 8-8-8 (Figure S5), and 22-8 (Figure S6), and one features a single barrel of 40 or more strands (Figure S7), larger than previously observed. Only in one MB-family the contact map contained clear signals that did not match our predicted architecture (Figure S9). That the predicted contact maps agree with the multi-barrel annotation predicted by the sequence homology methods, and that only one of the five clear contact maps conflicts with the homology-based predictions, suggests that most of the other architectures predicted only by sequence homology are also correct.

### Functional and taxonomic analysis of the multi-barrel MB-families

To better understand the significance of multi-barrel proteins and the evolutionary pathways that may have led to their emergence, we study their predicted functions and the taxonomic distribution within each MB-family. We predict the functions by homology-transfer, as identified by HHpred, and predict disordered regions using Quick2D [24]. In cases where multiple MB-architectures are evolutionarily linked, we used these relationships to trace their evolution. From the 186 MB-families, we elaborate on: two MB-families in which the functions of the individual barrel domains are complementary and an MB-family with representatives in *Escherichia coli* (*E. coli*).

#### 1. The PLA1-PLA1 (12-12) MB-family

MB-family 052 includes 6 proteins from γ-Proteobacteria with an architecture of 12-12 (Figure 1, right 12-12 marked with two stars). The proteins in this MB-family have two repeated 12-stranded OMBBs homologous to Outer Membrane Phospholipase A (OMPLA, annotated as PLA1 in Pfam), connected by a linker of 40 to 50 residues. OMBBs of the OMPLA MB-family are known to natively form homodimers, suggesting that in this case inter-*chain* interactions may have been replaced with inter-*domain* interactions. Possibly, the inter-domain interactions facilitate regulation.

OMPLAs are widespread in Gram-negative bacteria and were implicated in the virulence of some pathogenic species [25], acting as hydrolases that recognize and act on a broad spectrum of phospholipids. Their monomeric form is inactive, and only becomes active upon calcium-dependent homodimerization [8, 9]. Dimerization forms two binding pockets at the interface between the barrels, each harboring two calcium-binding and two active sites, which we label I and II. While residues from both monomers are involved in forming each calcium-binding site, all active residues in one active site belong to the same barrel. These residues form a catalytic triad in each active site, which has phospholipase A1 and A2, lysophospholipase A1 and A2, and mono− and diacyl glyceride lipase activities, hydrolyzing phospholipids to fatty acids and lysophospholipids [25].

The 052 MB-family includes 6 proteins, all belonging to γ-Proteobacteria. Four are from three *Oceanospirilalles* species (*Zymobacter palmae, Halotalea alkalienta* and *Carnimonas nigrificans*) and the remaining two belong to the *Agarivorans* genus (*A. albus* and *A. gilvus*, Figure S10). Proteins in this MB-family are within a conserved genomic environment only in *Oceanospirillales*, composed of genes encoding proteins that depend or act on cations. In *Agarivorans*, such an environment is absent (Figure S11).

When the individual barrel domains of the six multi-barrel proteins are clustered together with the barrel domains of single-barrel OMPLA proteins (Figure S12), the barrel domains from the multi-barrel proteins in *Oceanospirilalles* cluster closely with proteobacterial single-barrel proteins, while those from *Agarivorans* are far separated, substantiating that, although small, this MB-family is the result of two independent duplication events from two different single-barrel ancestors (Figure S10).

Figure 4 shows a homology model of a double-barrel based on the homodimeric structure of the *E. coli* single-barrel OMPLA as template. Sequence comparison suggests that all catalytic and calcium-binding residues are conserved in both barrels from *Oceanospirillales* but only in the second barrel from *Agarivorans* (Figure S13). In the first barrel from *Agarivorans*, two out of the three catalytic residues and three out of four calcium-binding residues involved in the formation of active site I are mutated to residues with biochemical properties incompatible with the function of their counterparts in *E. coli*, indicating that this active site was either lost or was able to evolve yet a different function. This latter observation suggests that one benefit of connecting the barrels within the same polypeptide chain may be the ability to diverge and explore functional adaptations not possible in a homodimer.

**Figure 4:**
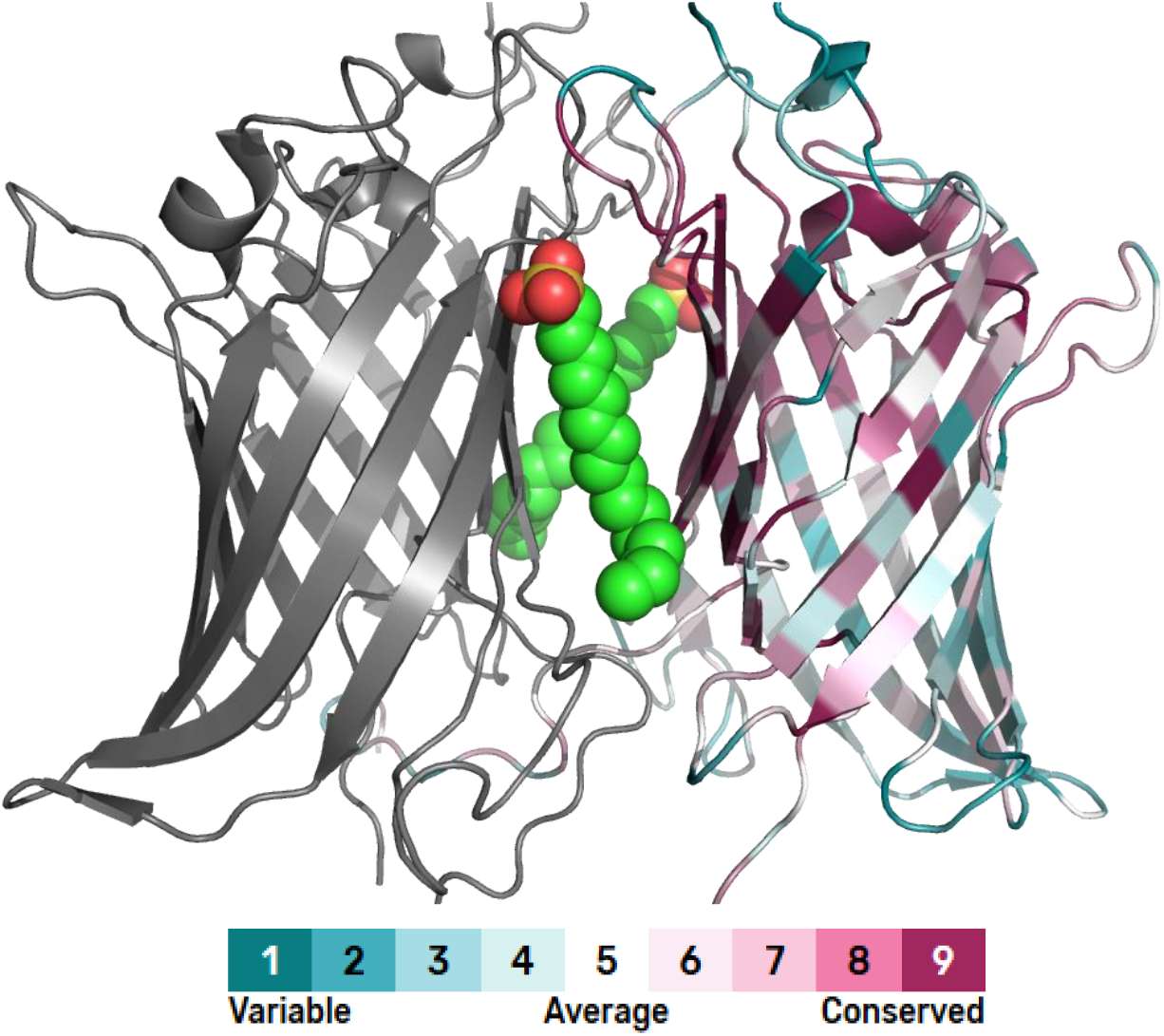
Homology model of a 12-12 double barrel based on the homodimeric structure of *E. coli* OMPLA (1QD5) [9] as template. One barrel is colored by the ConSurf conservation grades of the E. coli homolog, and the other in grey. Based on the template structure, two inhibitor molecules are tentatively docked in the interface between the barrels. The core of the barrel and the interface between the barrels are evolutionarily conserved, and the residues facing the membrane are variable.

#### 2. The PagP-LptD (8-26) MB-family

MB-family 093 is composed of 19 proteins from the Rhodocyclaceae branch of β-Proteobacteria with an 8-26 architecture (Figure 1, right 8-26 marked with a star), annotated as a combination of a Lipid A-modifying barrel of 8 strands (PagP) and the 26-stranded LptD barrel of the lipopolysaccharide transport (Lpt) complex, connected by a loop of more than 100 residues predicted to be disordered. Here, the two barrels share the same lipid substrate, and the connection between them might increase the efficiency of the process.

Lipid A is the lipid component of Lipopolysaccharide (LPS), the major component of the outer membrane of Gram-negative bacteria [26], which is synthesized in the inner membrane and transferred to the outer membrane by the LPS transport system. The last protein in the transport system is LptD, which transports LPS to the outer leaflet of the outer membrane. The link to the Lipid-A modifying barrel might shorten the distance an LPS molecule has to travel between the transport system and the modifying enzyme, thereby increasing the efficiency of the modification. This MB-family comprises proteins from β-proteobacteria, specifically species of the *Propionivibrio* and *Rhodocyclus* genera, whose conserved genomic environment is the same as that of LptD but not PagP in *E. coli* (Figure S14).

#### 3. The YjbH-GfcD (12-12) MB-family

This MB-family is the largest in our set, comprising 4,187 proteins, mostly from Proteobacteria (Figure 1, top-right 12-12 marked with two stars), with representatives also from other bacterial phyla. It is of particular interest as it is the only one with a representative in *E. coli.* Proteins from this MB-family can be divided into several subfamilies using CLANS with an E-value of 10^−5^. The central one encompasses 3,440 proteins (82%) from almost exclusively γ-Proteobacteria. The second largest is composed of 537 proteins (13%), almost exclusively from α-Proteobacteria. The third is composed of 82 proteins (2%), and has representatives mostly from α and γ-Proteobacteria. Two smaller subfamilies are originated from Chlamydiae (30 proteins) and from δ-Proteobacteria (26 proteins). The rest of the subfamilies are small, and contain mostly Proteobacteria. Five sequences in this MB-family are attributed to Euryarchaeota, a phylum in archaea. It is very unlikely that the sequences fold into functional barrels. Possible explanations are that the sequences are attributed to archaea due to sequencing error, or that they were horizontally transferred to archaea, but are non-functional.

Two paralogs of these multi-barrels, YjbH and GfcD/YmcA, are found in related operons in *E. coli* (Figure S15) and are clearly connected to the production of exopolysaccharide capsules [27–29]. Of these, the *gfcABCDE-etp-etk* operon has been studied extensively for its role in the formation of the group 4 capsule polysaccharide, and the structures and functions of most proteins encoded are understood (Figure S15) [30–33], except for GfcA (which is a short protein predicted to be natively unstructured) and GfcD. The yjbEFGH operon, which is a paralog of *gfcABCD*, has also been implicated in the production of an extracellular polysaccharide, but is much less studied. Both YjbH and GfcD are predicted to carry an N-terminal OMBB of the FapF family, a middle globular peptidoglycan-binding domain, and an unknown C-terminal OMBB without full-length homology to any known OMBB family, but predicted to be composed of 12 strands. The molecular advantage in coupling these two OMBB domains into one protein is unclear, as neither domain has close homologs among single barrels.

##### Poly-8 MB-Families

Finally, tandemly repeated 8-stranded OMBBs are the most common architectures in our set, with many different MB-families from diverse taxonomic groups emerging in parallel from different 8-stranded single barrels (Figure 1). Some of the MB-families can be grouped to two MB-superfamilies, containing evolutionarily related MB-families. The first poly-8 MB-superfamily includes 289 proteins from 28 MB-families, with between 2 and 11 OMBB domains, found only in Bacteroidetes (Figure 1, multiple architectures near the bottom marked with a star), especially in species from the *Prevotella* (105 proteins) and *Bacteroides* (157 proteins) genera, which represent the dominant bacteria of the human microbiome [34]. The second poly-8 MB-superfamily includes 118 proteins, with 2 or 3 barrel domains and an α-helical linker domain, found in various Bacteroidetes genera. The evolution of the two MB-superfamilies includes both amplification events that formed the (8)_11_ protein from an 8-8-8 ancestral protein, and deletion events that formed 8-8 proteins from 8-8-8 proteins, and (8)_6_ from (8)_7_ and (8)_9_ proteins. Fully tracing the evolutionary pathways that formed the poly-8 proteins can reveal insights to the evolutionary process itself and will be described in a separate manuscript.

## Conclusions

Given their internal symmetry and their distinction from soluble forms, OMBBs are an attractive class of folds for tracing evolutionary processes [6, 12, 35]. Scientists have long been familiar with proteins forming single OMBB domains [6, 12, 13], as well as with complexes of multiple barrels such as the OMPLA homodimer [8, 9], the porin trimer [10, 11], and the type-9 translocon heterodimer [7]. Our study allowed us to identify several novel OMBB forms, including the largest currently reported, and many proteins with multiple OMBB domains in the same chain, often in tandem arrangement. The existence of such multi-barrel proteins has not been established to date, although it has been a matter of conjecture [13].

In soluble proteins, the concatenation of two domains is expected to yield a two-domain protein. It is not obvious that this also applies to the membrane-embedded beta barrels, since, owing to their nature, their concatenation in tandem may also result in a single barrel of greater diameter, as shown with the concatenation of two 8-stranded OmpX barrels by Arnold et al. [14]. Indeed, Remmert et al. [12] and Franklin et al. [6] suggested that some large barrels evolved through the addition of β-hairpins to smaller barrels. In the cases we describe here, the multi-barrels almost invariably combine full-length matches to existing single-barrel proteins, connected by substantial linkers with a median length exceeding 20 residues. This is in contrast to the extremely short linker of 2 residues used by Arnold et al. [14] to successfully connect the two 8-stranded barrels into a larger barrel. Additionally, in the cases for which we could predict contact maps, these clearly support the presence of multiple independent barrels along the polypeptide chain.

The large number of multi-barrel architectures that we identified is notable in light of the conservative search procedures that we used. In particular, to minimize the number of falsely reported architectures, we (1) inspected only proteins with at least two full matches to the seed sequences; and (2) inspected the proteins we found with HHpred and further sequence annotation tools. We believe that with a larger set of seed sequences, or using more sensitive homology search tools, it is likely that we would have found even more multi-barrel architectures. Even with these conservative choices, we found many multi-barrels from a multitude of bacterial phyla, highlighting how frequent these proteins are in Nature.

Our example analyses show that multiple evolutionary processes lead to the diversity of multi-barrel proteins. One is descent with modification from an ancestor that already contained multiple barrel domains, as illustrated by the YjbH/GfcD family, which is ancient and conserved in many bacterial lineages that have an outer membrane. Another is the amplification of a beta barrel that normally forms oligomers for functionality into a single chain with multiple barrel domains, as observed in two independent lineages of outer membrane phospholipase A. Also, we saw the fusion of different barrels acting within the same biological process, as described for the PagP-LptD MB-family involved in lipopolysaccharide biogenesis.

The poly-8 MB-superfamilies allow one to trace the evolutionary events forming them. A preliminary examination revealed lineage-specific deletion of individual domains from a multi-barrel ancestor and multiple independent amplification of barrel domains. A detailed study tracing the evolutionary relationships in among these multi-barrels is underway. The diversity of these evolutionary mechanisms underscores the frequency with which multi-barrels occur in many lineages.

We find that multi-barrels evolve frequently, but only some of these appear to be retained over long evolutionary periods. Most appear in only one or a few bacterial genera and some even only in individual species, suggesting a large turnover through *de novo* evolution and subsequent gene loss. This is in agreement with the results of genetic processes in general, which typically remain lineage-specific even when they become fixated in the genome and disappear when the lineage dies out. Nevertheless, the large number of multi-barrels that have become fixated in genomes, even only with a very narrow phylogenetic distribution, begs the question of the biological advantages they may confer over their single-barrel homologs. Several can be envisaged: (1) increasing the efficiency of a given pathway, as we propose for the PagP-LptD fusion in lipopolysaccharide biogenesis; (2) opening avenues for the divergence and separate functional adaptation of domains previously engaged in homo-oligomerization, as in the outer membrane phospholipase A; and (3) increasing the avidity of proteins engaged in binding multiple adjacent epitopes. This last consideration may offer a hypothesis for the prevalence of poly-8 proteins in the gut genera *Bacteroides* and *Prevotella*, since one of the functions reported for OmpA is adhesion [36] and the long polysaccharides in the human gut offer many adjacent epitopes of the same or similar nature.

An important open question in understanding multi-barrels is their mechanism of insertion in the outer membrane. In single-barrel outer membrane proteins, this process is mediated by the β-barrel assembly machinery (BAM) complex [37], which recognizes a sequence signal in the C-terminal strand of the barrel to be inserted. The process has been studied mainly in Proteobacteria and mitochondria, so the nature of the sorting signal is still understood only in general terms and may be partly lineage-specific [38–40]. In some of our multi-barrels, mainly in those that have clearly arisen recently, we can readily recognize the presence of the sorting signal in each constituent barrel, but this may be the result of recent fusion, as opposed to the need for two independent signals within the same protein. So far, we have not attempted to build a prediction tool for sorting signals and are not aware that such a tool is available. Hence, it is unknown if one signal at the C-terminus is sufficient for the insertion of the entire protein into the outer membrane, or each barrel is inserted independently by the BAM machinery.

The extent of OMBBs identified herein highlights their functional importance. Furthermore, OMBBs may have even more undescribed functions: a preliminary search that we carried out, revealed many proteins combining beta barrels and non-barrel domains; the latter include DNA binders, histidine kinase sensors, and ABC transporters. Overall, the novel architectures identified herein suggest that OMBBs have an even greater functional role than was previously thought.

## Methods

We used two methods to gather protein sequences likely to have multiple barrel domains: A HMMER-based method and a PsiBLAST-based method. The sequences found by either method were clustered by similarity, and the clusters were manually inspected to ensure every cluster had a single architecture (Figure 5).

**Figure 5:**
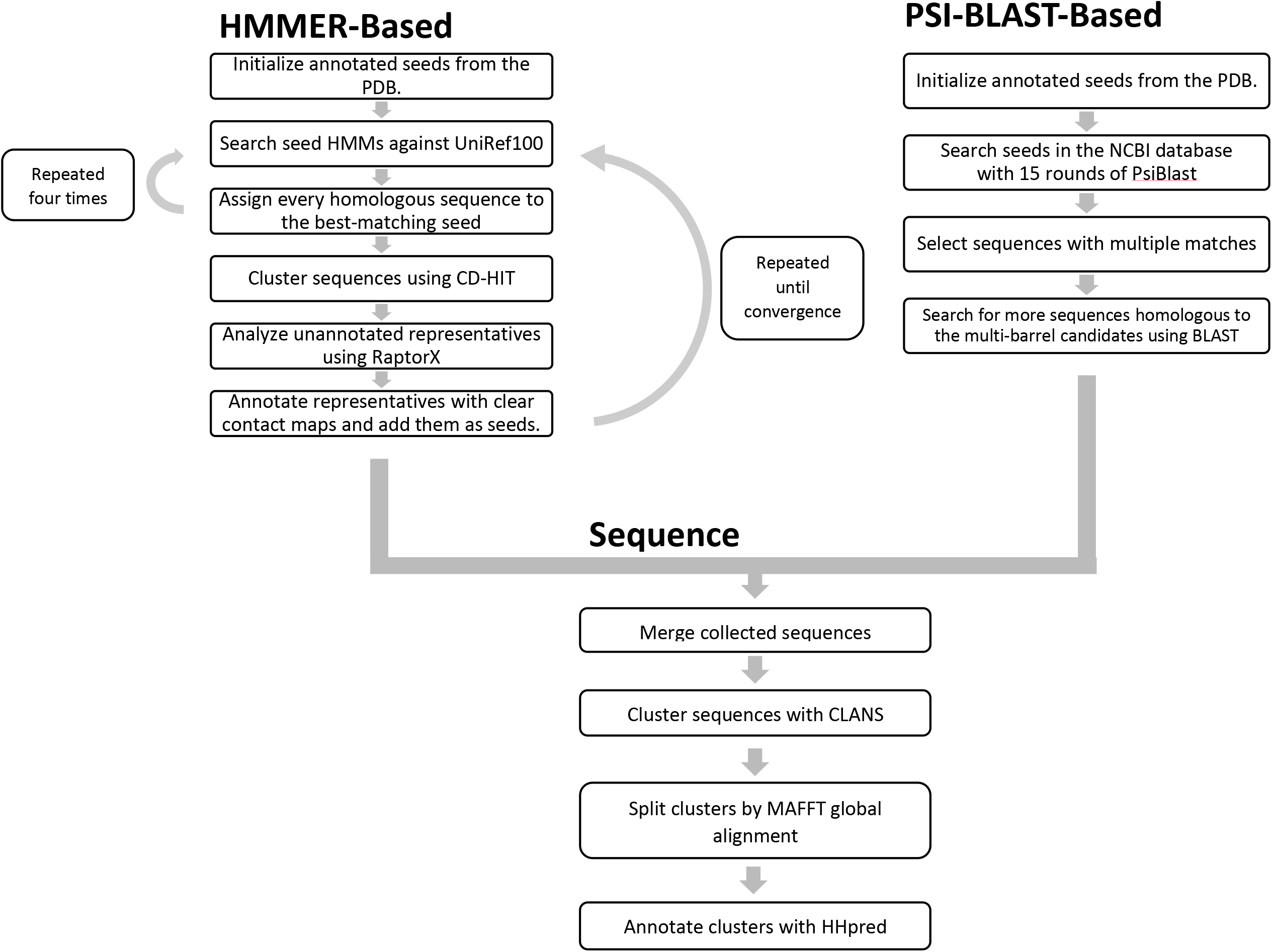
A flowchart of our protocol to find multi-barrel proteins.

### HMMER-based sequence searches over UniRef100

An overview of the search procedure is depicted in Figure 5, and the steps are described below.

#### 1. Constructing the set of seeds (Single-barrel HMMs annotated with barrel size)

We started with all 275 PDB sequences of OMBB proteins as identified by MemProtMD [41]; the most recently solved structure in the set we used is 6FSU (released 11/2018). We trimmed the N− and C-terminal ends that were not part of the barrels. Then, we selected from this set a less-redundant subset of 98 proteins using CD-HIT v4.7 [42] and a 90% identity threshold. Using the corresponding structures, we annotated each protein sequence with the number of strands in its barrel.

#### 2. Enriching the seed HMMs

For each OMBB in our set, we identified homologs iteratively, building our seed HMMs in four rounds. Initially, the set of homologs for each OMBB was the trimmed sequence itself. In every iteration, we used HMMER 3.1b2 [17] to list all the sequence homologs in UniRef100 of any of the seed HMMs (using an E-value threshold of 10^−5^). Then, we traversed the list in ascending order of E-value: for every sub-sequence that matched a seed, provided it did not intersect with already-considered subsequences on the same sequence, we added it (only) to its best-matched seed. This way, a sequence with high similarity to one barrel and low similarity to another would be added only to the set of homologs of the former. We further filtered the set of homologs, enforcing high coverage: the sequences must match 90% of the seed HMM. Finally, we added the identified homologous sections to the seed alignment and rebuilt the MSAs and HMMs of the seeds. This procedure constructed seed HMMs that included remote homologs from the same family, while not being corrupted by close homologs from a different barrel family.

#### 3. Adding seeds for new predicted single OMBBs

In the process of building the HMMs for the seeds, we found UniRef100 sequences that could not be added to the set of homologs of any seed, because the coverage of their matches was too low. These sequences may be members of OMBB families that are not represented in the PDB, or remote homologs of families that are represented. Given that we wanted to find as many OMBB families as possible, we inspected representatives of these sequences, and added those which we found to be OMBBs to the seed set. We performed four rounds of new seed selection, and after each round we reconstructed the HMMs using the same protocol described above.

To limit the number of representatives checked, we selected only the subset that is more likely to contain barrel domains. For that, in the first three rounds of new seed selection, we collected sequences with at least two partial matches. We clustered them with CD-HIT and used RaptorX to construct an MSA and to predict a contact map for a representative from each cluster. When the predicted contact map of a sequence had a clear β-barrel pattern from which we could infer the strand number via a manual inspection, we concluded that this is likely an OMBB and added the barrel domain as a seed. In the fourth and last round, we collected all sequences with only a single partial match, clustered them using CD-HIT, and picked representatives from the largest clusters. We predicted contact maps for the representatives using RaptorX, and if the predicted contact map manifested a clear β-barrel-like pattern we added them as seeds as well. Our process converged after the third and fourth rounds of new seed selection.

#### 4. Using the seed HMMs to identify multi-barrel proteins

Our goal was to find with high sensitivity multi-barrel proteins, and to accurately annotate their individual domains. To achieve this, we started with a two-step procedure to collect a set of multi-barrels, aimed at minimizing the false-negatives: (1) we searched UniRef100 for homologs of the proteins in our seed set, using HMMER and the very lax E-value of 1000. This was the initial database of potential homologs, which were possibly false. (2) Within this initial database, we searched again using a stricter (maximal) E-value of 0.1.

The challenge in annotating the barrel domains of the sequences found in UniRef100 is that each sequence has many alignments to our seeds, sometimes matching the same residues in that sequence. To address this challenge, we devised the following procedure: First, we identified and corrected false local matches. HMMER searches for local alignments between the seed HMM and the UniRef100 sequences. Because of this, large insertions split full-length matches between the seed HMM and the target sequence into multiple non-overlapping short matches. To correct this, our program merged these fragments into a larger single match. Then, we used a greedy approach to iteratively collect a list of the trusted matches for each target UniRef100 sequence. Initially, the trusted list was empty. We traversed the list of all the target sequence matches to the seed HMMs in ascending order of E-value. For every match, we checked if it had long intersections with matches that were already in the trusted list. If there were intersections longer than 10% of the shorter match, we discarded the currently considered match. Otherwise, we added the match to the list of trusted matches. Finally, relying on this list, we used the barrel sizes of the matching seeds to annotate the sequence.

Having collected and annotated the set of candidate sequences, we removed from our set sequences with fewer than two full matches, or with fewer than three partial matches; we then clustered the sequences using CLANS [43] (and an E-value of 10^−20^). In cases where the cluster had sequences annotated with different sizes (e.g., a cluster with sequences annotated with two consecutive 8-stranded barrels, and sequences annotated with an 8-stranded barrel followed by a 10-stranded barrel), we further split the cluster, so that all sequences in each cluster would have the same barrel annotation. We used MAFFT v7.409 [44] to build an MSA for every cluster, and HMMER to expand these MSAs four times.

The search used 239 seed HMMs. In the initial search, 1,387,337 proteins with at least one partial match to the seed HMMs were found. After removing proteins with fewer than two full matches or three partial matches, 4,194 proteins which were likely multi-barrels were found.

The 4,194 proteins were divided to clusters, and the clusters were divided by architectures. For every architecture in every cluster we built an HMM and performed another HMMER search against UniRef100. After this final search, we had 9,062 annotated multi-barrel proteins.

### PsiBlast-based sequence searches over NCBI’s non-redundant protein sequences database (*nr*)

Independently, we used another approach to find proteins with multi-barrel matches that complements the method described below (Figure 5, upper-right).

#### 1. Constructing the set of initial sequences

To collect a set of reliable initial OMBB protein sequences, global and local structural homologs of 6 OMBB families were collected using HHpred. Input sequences corresponded to *E. coli* BamA (UniprotKB P0A942), *E. coli* FepA (UniproKB P05825), *E. coli* OmpX (UniprotKB P0A917), *E. coli* Phospholipase A (UniprotKB P0A921), *K. pneumoniae* LptD (UniproKB C4T9I0), and mouse VDAC1 (UniprotKB Q60932). Searches were carried out over the PDB database filtered to a sequence identity of 70% as of January 2019, using default parameters. The resulting PDB entries matched were collected, the barrel identified, its sequence saved along with the number of strands, the length of internal and external loops collected and any N- and C-terminal decorations trimmed. In total, 157 unique OMBB sequences were collected, with barrel topologies ranging between 8 and 36 strands.

#### 2. PsiBlast searches for bacterial proteins with multi-barrel matches

These OMBB sequences were then used as seeds for PsiBlast searches over the bacterial sequences in the NCBI’s non-redundant sequence database, filtered to a sequence identity of 70% (nr_bac70). For all seeds, a maximum of 15 PsiBlast rounds were carried out, using those matches with an E-value better than 10^−4^ to generate the PSSM for the next round, and saving the hits with an E-value better than 10 after each round. The searches were carried out with PsiBlast 2.6.0+, allowing for the filtering out of low complexity matches with SEG (-seg option on). All matches with a minimum query coverage of 90% were then processed together, to find sequences with at least 2 non-overlapping matches to any of the queries. Given that domain boundaries are always difficult to define with certainty at the sequence level, two matches were considered non-overlapping even if they overlapped by up to 10 residues. This search resulted in 482 unique EntrezIDs comprising multiple OMBB matches in a single chain. As we observed that matches to the large 36-stranded OMBB were generating ambiguous matches to families with a well-defined barrel topology (e.g., 22-stranded barrels of the FhuA family), these matches were excluded from further consideration.

#### 3. Enriching the set of bacterial multi-barrel proteins

To enrich this set of proteins containing multi-barrel matches, we used the 482 unique, full-length, sequences as seeds for a BLAST search over the complete set of bacterial NCBI’s non-redundant sequences (nr_bac). Searches were carried out as described above, collecting matches with an E-value better than 1 and a sequence coverage of the multi-barrel region higher than 90%. The resulting set of full-length sequences was then filtered to a sequence identity of 100%, eliminating any redundant match. A total of 6992 sequences were collected. Here, no preliminary topology annotation of putative multi-barrel sequences was carried out based on the queries as the deep searches preformed in step 2 may lead to the identification of larger barrel topologies when starting from a smaller number of strands (e.g., due to indels).

### Classification and domain annotation

To perform a rough clustering of the sequences and visually inspect the clusters, we used CLANS [43]. CLANS takes a list of protein sequences as an input, performs all-against-all BLAST comparisons between the sequences, and stores the p-value of the similarity of every similar sequence pair. To allow visual inspection of the sequence clusters, CLANS represents every sequence as a point in a two dimensional space and places the points in a way that minimizes the distance between points representing homologous sequences, while keeping points representing non-homologous sequences away from each other.

To allow a finer clustering of similar sequences, CLANS can use a p-value cutoff, and consider only distances between sequences whose homology has a p-value smaller than this cutoff.

The proteins collected using both the HMMER-based and a PsiBLAST-based approaches were combined, redundant sequences excluded with CD-HIT at a sequence identity of 100% and clustered with CLANS [43]. Clusters were identified at a p-value of 1×10^−25^ and only those comprising at least 3 sequences were considered. The p-value was picked manually so that each MB-family will be clustered to a single cluster, but different MB-families may also be clustered together.

To assign to every protein its architecture, we split the clusters found by CLANS further. For every CLANS cluster, we calculated an MSA using MAFFT. Then, we split the proteins to sub-clusters using the alignment. Two proteins were determined to be in the same sub-cluster if they shared more than 70% of their positions in the MSA. This made sure that proteins with a completely different architecture would be in different sub-clusters, since the unmatched barrels would create a large number of unmatched positions in the MSA. This method ensures that the proteins in every cluster are aligned to each other well for further analysis.

After splitting to sub-clusters, we separated close architectures. Two architectures could be sent to the same sub-cluster if one was contained in the other – a protein with an 8-8 architecture can share most of its positions with an 8-8-8 architecture protein. To separate those, we manually inspected the distribution of the lengths of sequences in the clusters. When there were sequences whose lengths were at least 100 amino acids longer or shorter than the most common length, we found their architecture using HHpred and checked if it differed from the architecture of the other sequences in the cluster. This method identified one cluster which contained false positives, and five small groups of sequences which were in the same cluster with sequences with different architectures.

This method also over-splits clusters when there are large insertions, or non-barrel domains. This caused cluster 15, which contains a large single barrel and has both long insertions and domains other than the barrel domain, to be split to many small clusters. To separate the proteins in cluster 15 to subclusters properly, we used two steps: We first identified the barrel domains, and then we applied the previously described splitting method using only the barrel domain in every sequence.

To detect the barrel domains automatically in the sequence, we used HMMs. We manually identified the barrel domains in six sequences using HHpred. We used them to create six initial HMMs, and for each HMM, we searched for homologs among the sequences of cluster 15, which we then added to the HMM. We repeated this process four times, until the barrel domains were detected in all proteins of cluster 15. We then split the proteins in cluster 15 using the similarity of the barrel domains. The cluster was split to four sub-clusters this way.

From every sub-cluster, a representative protein was analyzed with HHpred, and the closest homologs were used to determine its architecture. When there were only partial overlapping matches to homologous proteins, we assumed the section was a single barrel, and determined its number of strands by HHpred’s secondary structure prediction and by the matches to strands of known structures.

At the end of the procedure, we had 186 sub-clusters, to which we refer in the paper as MB-families.

### Homologous structures and contact prediction

Our MSAs include multi-barrel proteins and other barrel architectures that extend the currently documented repertoire. We used sequence analysis tools to obtain further support for the correctness of our predicted architectures. First, we used HHpred [45] as an alternative homology-based tool to remove likely erroneous cases. To this end, we constructed and searched for HMMs (one round of PSI-BLAST, no secondary structure scoring) in the PDB, ECOD, and Pfam. This procedure identified five cases that were predicted by HHpred to have only a single barrel domain, and we removed them from our datasets.

We also used the RaptorX webserver to predict contact maps. Because contact map prediction relies on the coevolutionary signal, it requires many homologous proteins. Thus, we could use the server to predict the maps only for the 23 families containing at least 50 proteins. RaptorX limits the length of the MSAs to 1300, and the MSAs of our families were often longer. To overcome the length limitation, we arbitrarily picked a single protein as query, and removed all positions in the MSA in which the query protein had a gap. Finally, we inspected each of the predicted contact maps to deduce a structural characterization of the multi-barrels. In many cases the β-barrel signal in the contact-map was missing, blurry, or riddled with too many false positives to use. We obtained a clear multi-barrel signal for only four cases out of 23.

### Structural modelling the PLA1-PLA1 (12-12) MB-family

To model the structure of the PLA1-PLA1 (12-12) double-barrel, we used HHpred to find structural templates for both OMPLA domains, and picked 1QD6, a homo-dimer bound to a hexadecanesulfonyl fluoride inhibitor. We cut the protein to two parts according to HHpred and removed the linker section, which was unaligned to the template.

To align the double-barrel sequences to 1QD6, we built an HMM for the two domains of one of the double-barrel proteins using HMMER. We searched for homologs in UniRef100 using an E-value of 10^−10^. We used MAFFT to create an MSA which contained the sequences of the two domains of the double-barrel, the sequence of the template, and sequences of other homologs of the 12-stranded barrel. We deleted all sequences in the MSA beyond the sequences of query and template and removed empty columns in the alignment. The result was an alignment of the domains of the double barrel to the template sequence, which used the information in other homologs of the OMPLA barrel to improve the accuracy.

We used this alignment to create a PIR file for MODELLER 9.23, and generated five models. We aligned the models we generated to the model of 1QD6, and copied the inhibitor from 1QD6 to the models we generated. We picked the model manually, attempting to minimize the steric clashes with the inhibitor.

We calculated conservation scores for the fused protein and for the template using ConSurf with the default settings. The N-terminal barrel domain of the 12-12 protein had 1,830 homologs, 150 of which were used. Since the PLA1-PLA1 (12-12) query has only six homologs, ConSurf calculated the conservation scores using homologs of the monomer and provides only an illustration of the conservation scores of the monomer and a sanity check for the predicted model.

### Classification of individual OMBBs from selected families and analysis of the genomic context

To assess the likelihood of the OMBBs in the PLA1-PLA1 MB-family to have emerged by the tandem repeat of an ancestral OMBB, we further classified the individual barrel domains with their single-barrel homologs in close species. For that, the individual barrel domains in the reference sequences selected for the respective family were used as seeds for a BLAST search over the nr_bac database, and only hits covering a minimum of 90% of the query collected. For the PLA1 proteins, given their high sequence identity to the single-barrel PLA1, only hits with an E-value better than 1×10^−40^ were collected in order to reduce the number of sequences to cluster. The resulting sets of individual barrel proteins were clustered with CLANS [43] with a p-value threshold of 1×10^−80^.

To assess the conservation of the genomic context of the PagP-LptD, PLA1-PLA1, YjbH-GfcD MB-families, the genomic context of each protein in the MB-family (or a set of representative cases) was carried out by identifying the products of the *n* flanking genes in their full genomic assembly and their further comparison using BLAST. For each case, clusters of homologous flanking genes were identified by all-against-all BLAST searches of their protein products at a maximum E-value of 1×10^−3^. Protein products from genes conserved throughout the genomic contexts analyzed were further annotated for their domain composition, secondary structure and sequence features as described above.

In the BLAST search of the PagP-LptD family, 17 additional proteins belonging to the MB-family were found.

### Calculating the taxonomic tree of the species containing multi-barrels

To display the taxonomic classification of the bacteria containing the multi-barrel proteins, we constructed a tree using NCBI’s taxonomy database [46], and rendered it using Dendroscope [1].

## Acknowledgements

This research has been supported by grant 94747 by the Volkswagen Foundation. NB-T’s research is supported in part by the Abraham E. Kazan Chair in Structural Biology, Tel Aviv University.

