## Supplemental Data for "Gram-negative outer membrane proteins with multiple β-barrel domains"

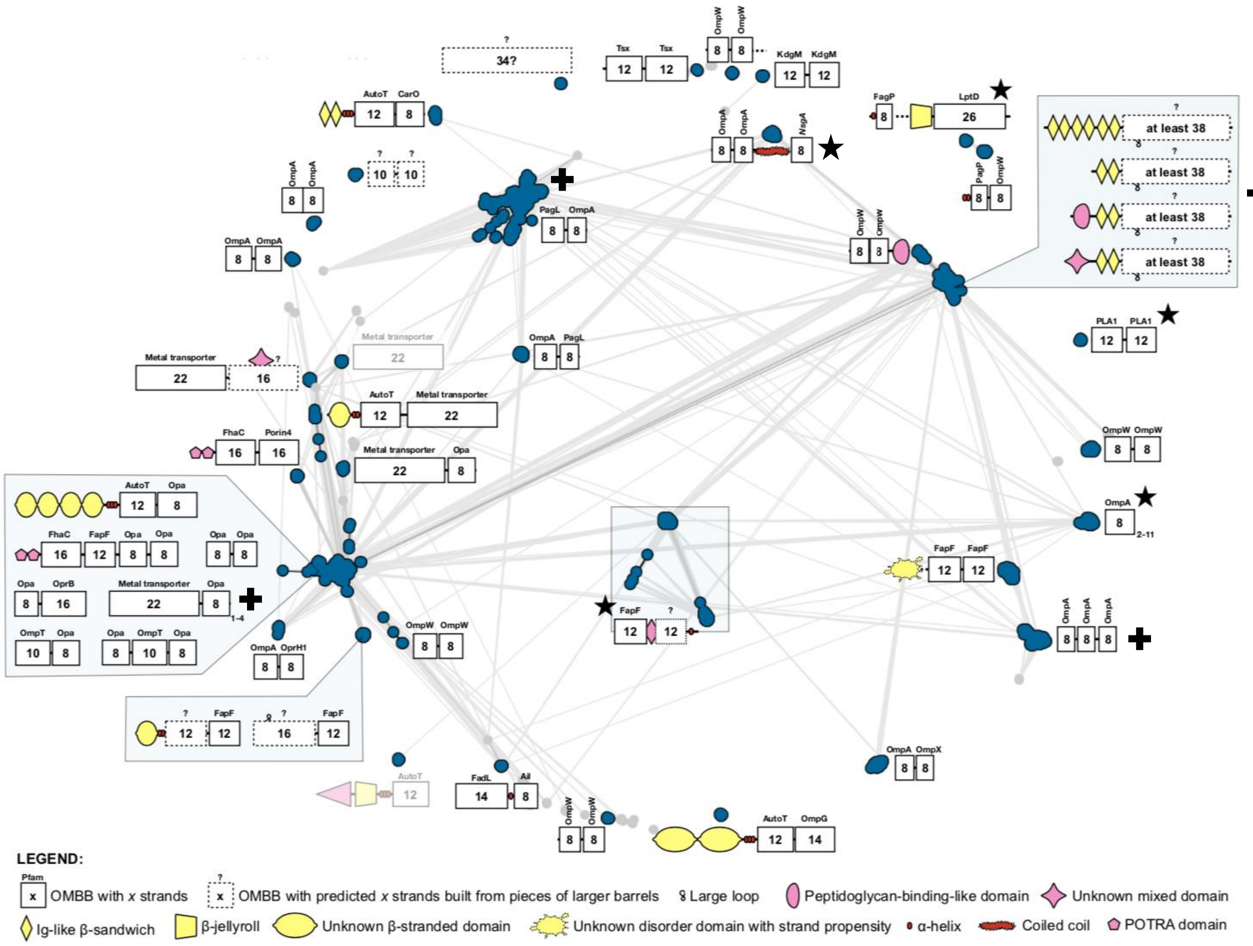


Figure S1. Classification and domain annotation of 12,643 proteins. Clustering was carried out with CLANS [1] at a p-value of $1\times10^{-25}$ and shown at $1\times10^{-15}$. For each cluster, sequences were binned by their size, with a step of 100 residues, and a representative from each bin collected. Each of these representatives was manually annotated for their predicted domain composition with HHpred [2] over Pfam [3], ECOD [4, 5] and PDB70 [6], for their secondary structure content with Quick2D [2], and for their transmembrane topology with BOCTOPUS [7]. Coiled coil domains were identified with PCOILS [8] and signal peptides with SignalP 5.0 [9]. Shaded architectures represent clusters that are composed of false-positive sequences carrying only one barrel belonging to a known OMBB family. Clusters with 3 or less sequences were not considered and are colored light grey. Black crosses mark four architectures with clear contact maps. Black stars mark the five families/MB-superfamilies selected for detailed discussion: The 12-12 OMPLA family (right edge), the 8-26 PagP-LptD family (top-right), the poly-8 MB-superfamily (bottom-right, marked with a single box), the 8-8-Linker-8 MB-superfamily (center-top), and the 12-12 YjbHGfcD family (center).

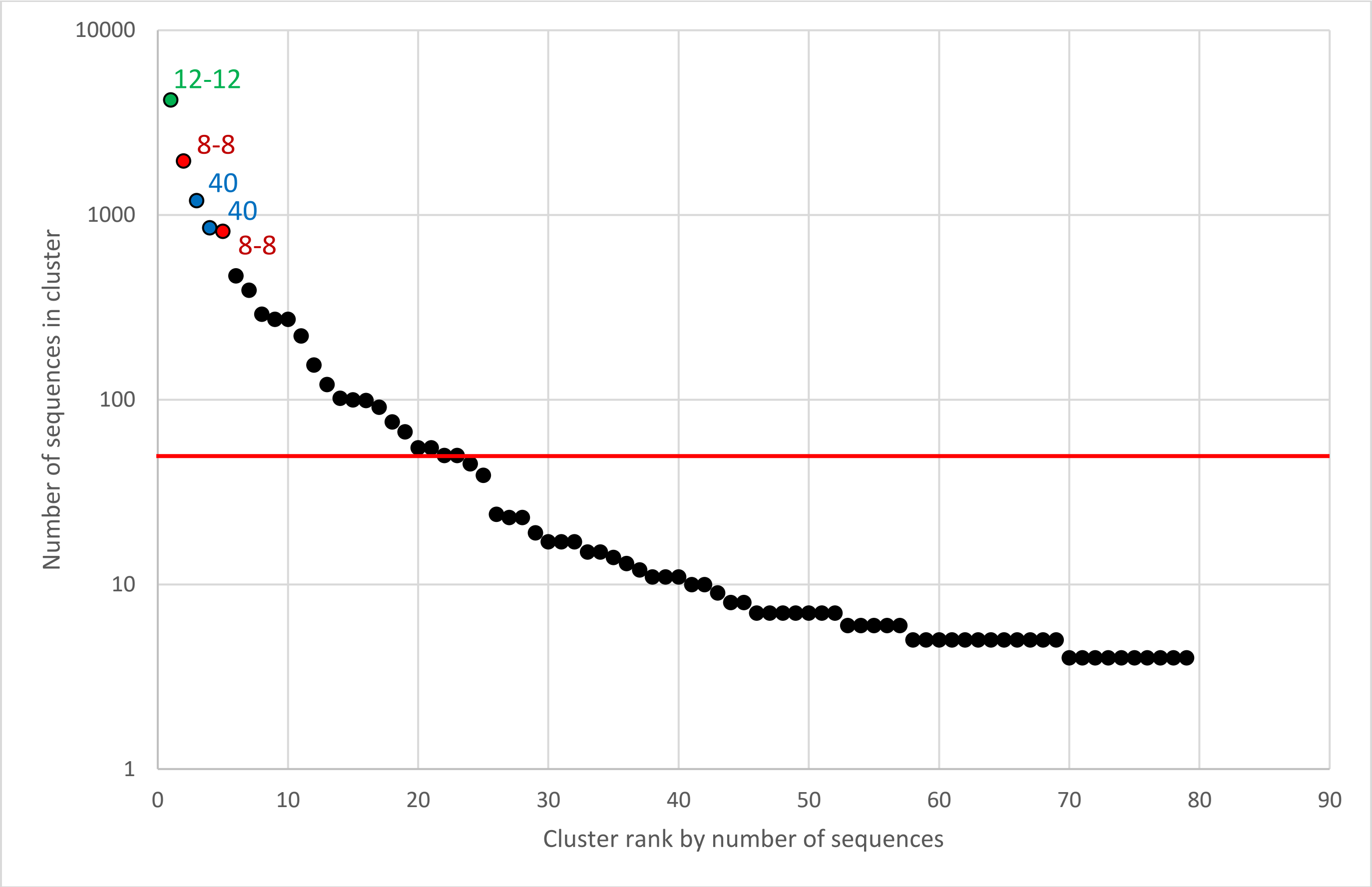


Figure S2. Families of barrels with new architectures (multi-barrels and large barrels) with at least four proteins, sorted by decreasing size, i.e., number of proteins per family, displayed in log scale. The points representing the five largest families are labeled and colored according to their MB-architecture – the 12-12 architecture in green, the 8-8 architecture in red, and the 40 architecture in blue. We see that quite a few MB-architectures are populated by many homologous proteins. The red line separates families with 50 homologous proteins or more, to which contact maps were predicted using RaptorX, and families with fewer than 50 proteins.

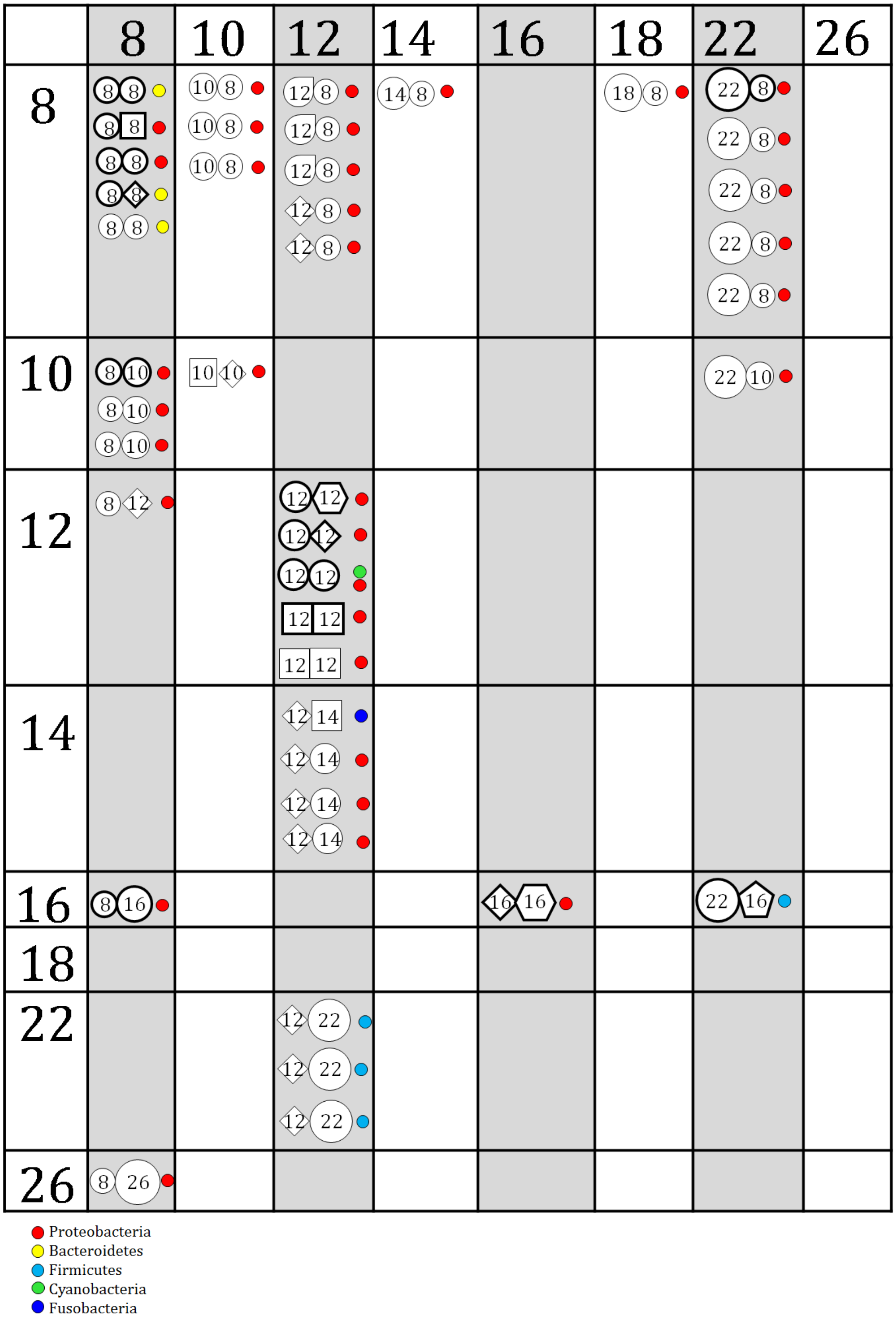


Figure S3. All pair architectures. The largest MB-families (up to five) within each architecture are shown. MB-Families with 50 homologous proteins or more are shown with a bold outline. The colored dot(s) show the taxonomy of the bacteria with the architecture. The topmost 12-12 architecture and the 8-26 architecture are discussed in the main text.

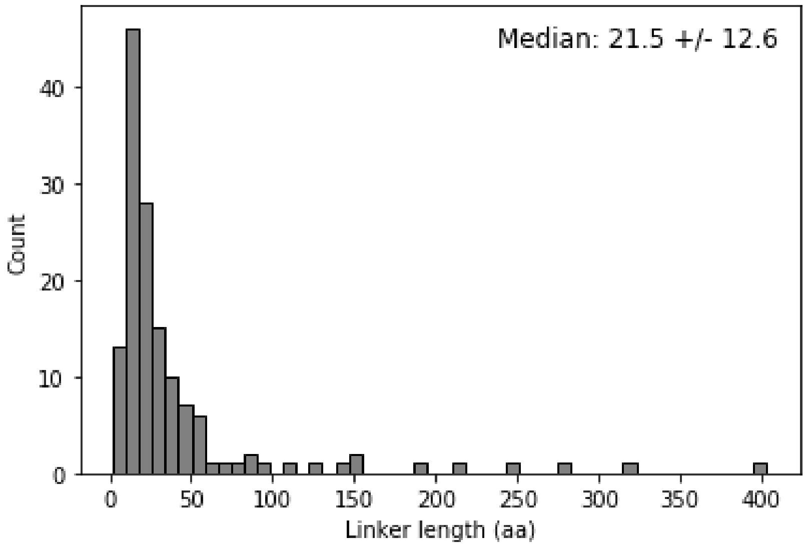

Figure S4. Histogram of lengths for the linkers between the barrels in the MB-families of Figure 1.

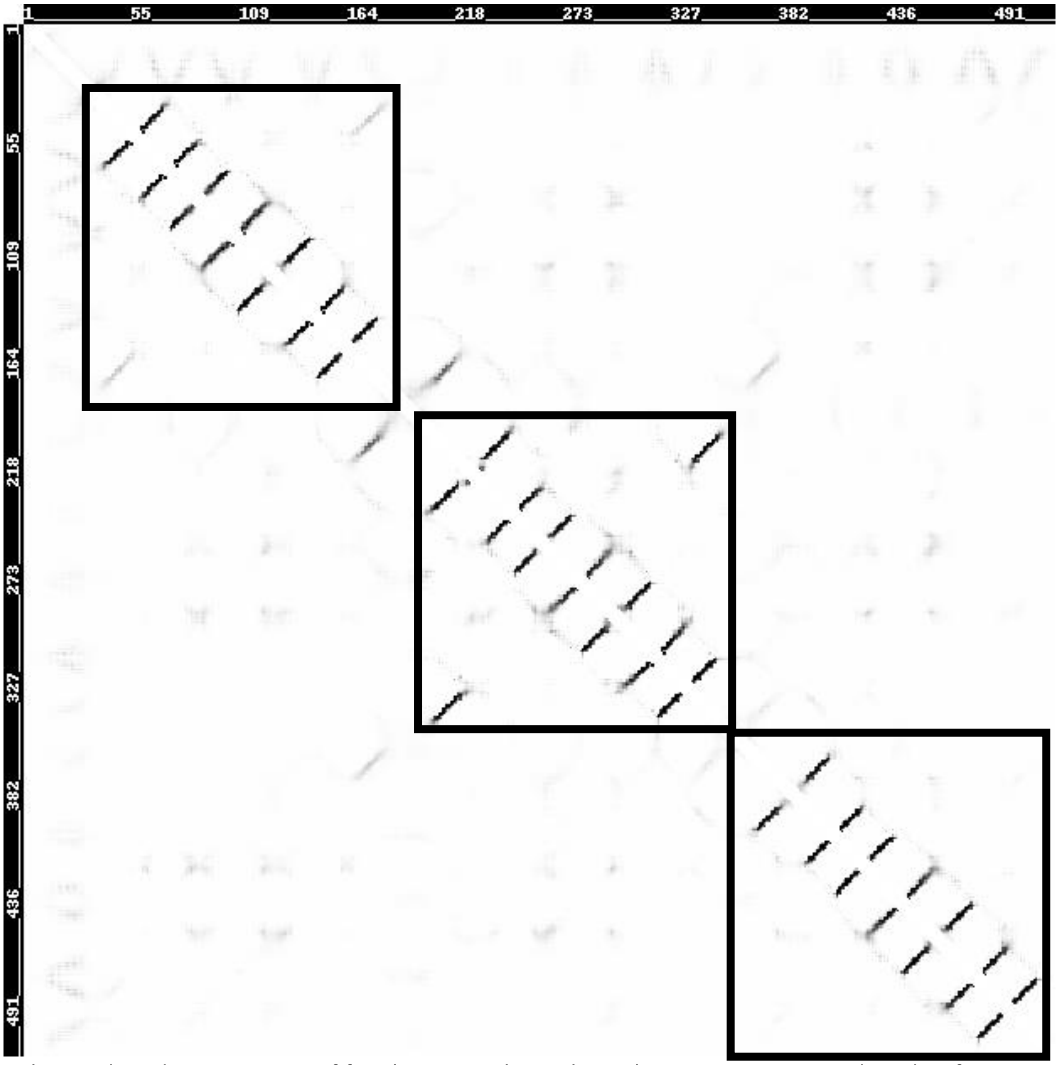


Figure S5. The predicted contact map of family 007, with 291 homologous proteins, predicted to form an 8-8-8 architecture. Clear contact signals are observed for a total of 24 beta-strands. Reassuringly, a clear 'barrel-closing signal', i.e., the contact between the first and last strands of the barrel, is observed for the middle barrel, and weak barrel-closing signal for the first barrel. A barrel-closing signal for the C-terminal barrel is missing. The three putative barrel domains are marked with black squares.

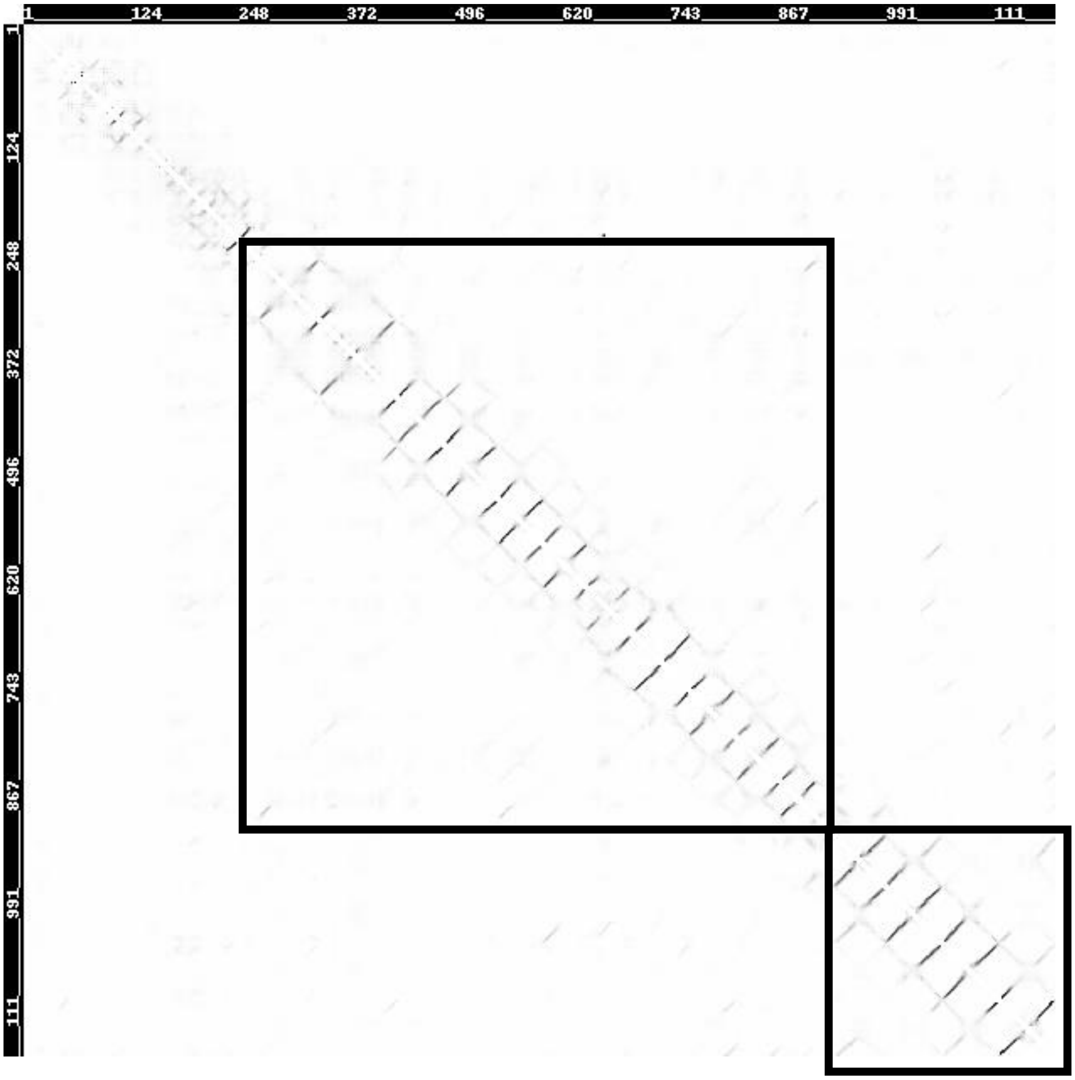


Figure S6. The predicted contact map of family 015, with 99 homologous proteins, predicted to form a 22-8 architecture. Clear contact signals are observed for a total of 30 beta-strands. Reassuringly, clear 'barrel-closing signals', i.e., the contact between the first and last strands of the barrel, are observed for both barrels. Furthermore, a barrel-closing signal between the stands that would indicate a single 30-stranded barrel is missing, making this alternative architecture much less likely. The two putative barrel domains are marked with black squares.

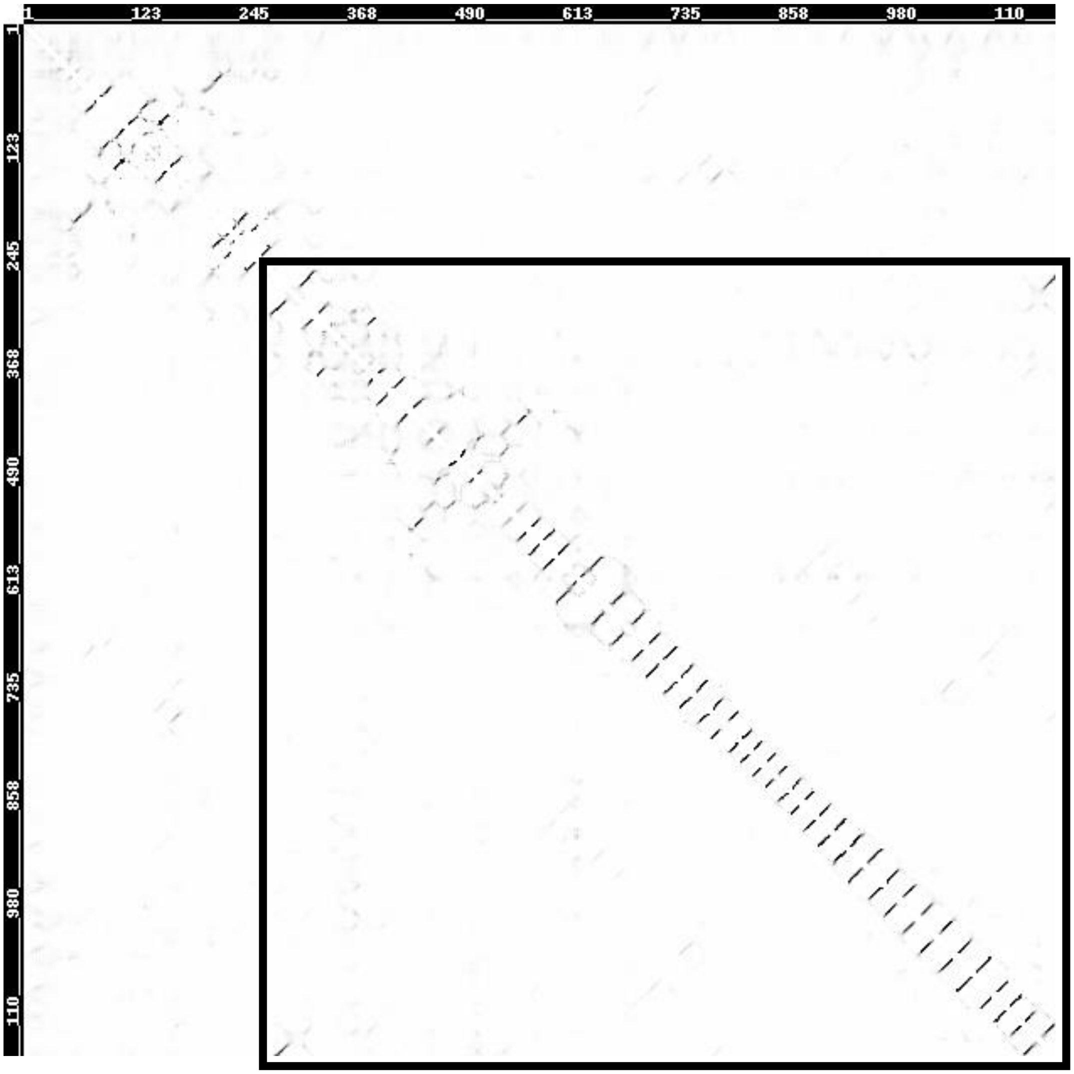


Figure S7. The predicted contact map of family 002, with 1,194 homologous proteins, predicted to form a single large barrel. Clear contact signals between adjacent strands are observed for a total of at least 40 beta-strands. Furthermore, a clear ‘barrel-closing signal’ between the first and last strands of the barrel is also visible. The putative barrel is marked with a black square.

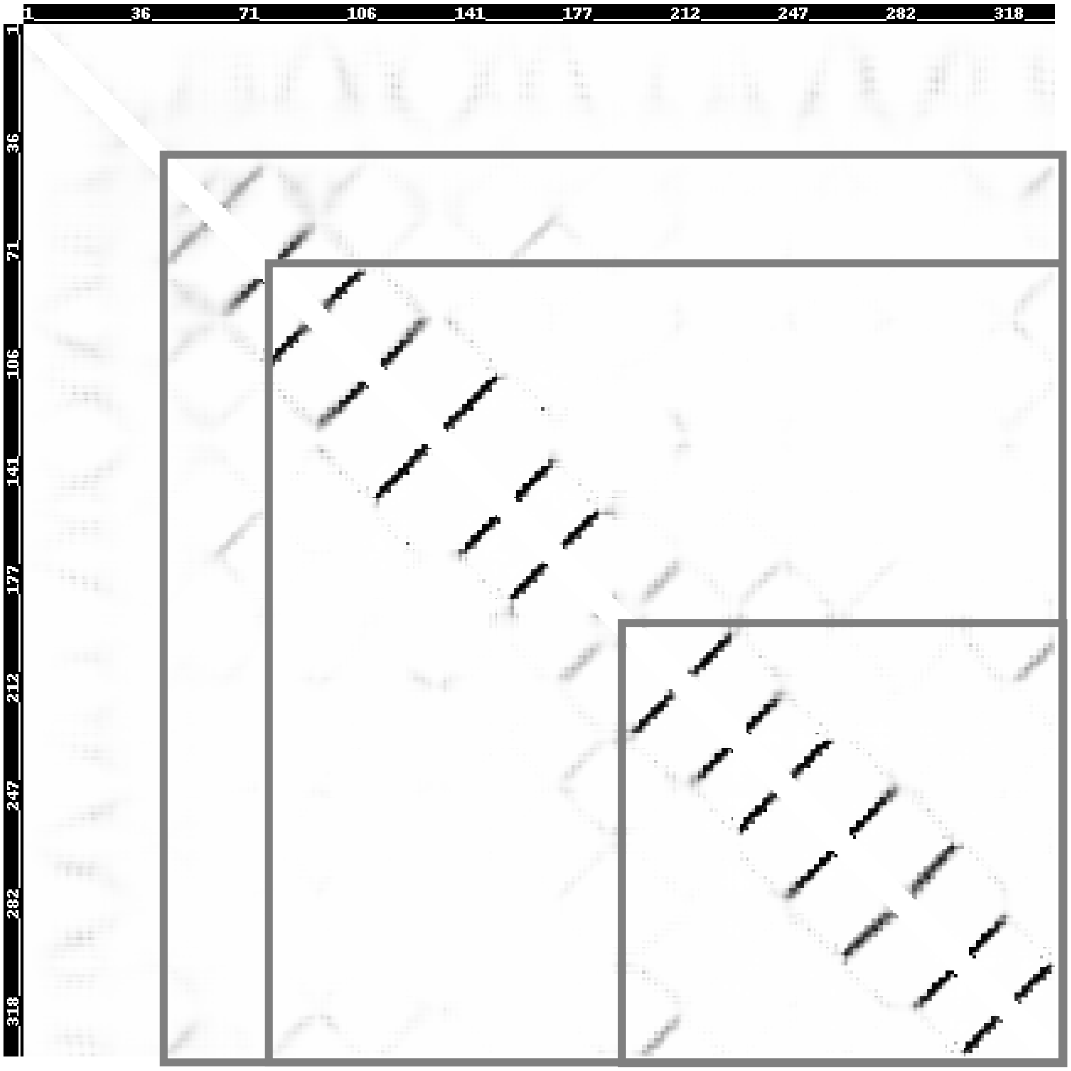


Figure S8. Example of an uninterpretable contact map: family 003, with 813 homologous proteins, predicted to form 8-8 architecture. The signals of contacts between adjacent strands within both putative barrels are clear. There is also a barrel closing signal for the second domain, in supports of the predicted 8-8 architecture. However, there is also a weak contact signal between the last strand of the first putative barrel domain and the first strand of the second putative barrel domain, suggesting a single barrel architecture. Furthermore, there are weak closing signals for hypothetical single barrel architectures with 14 and 16- strands, as well a weak closing signal between the second strand of the first barrel and the seventh strand of the first barrel, which is unlikely. Overall, the contact map does not provide unequivocal evidence in support of a single architecture.

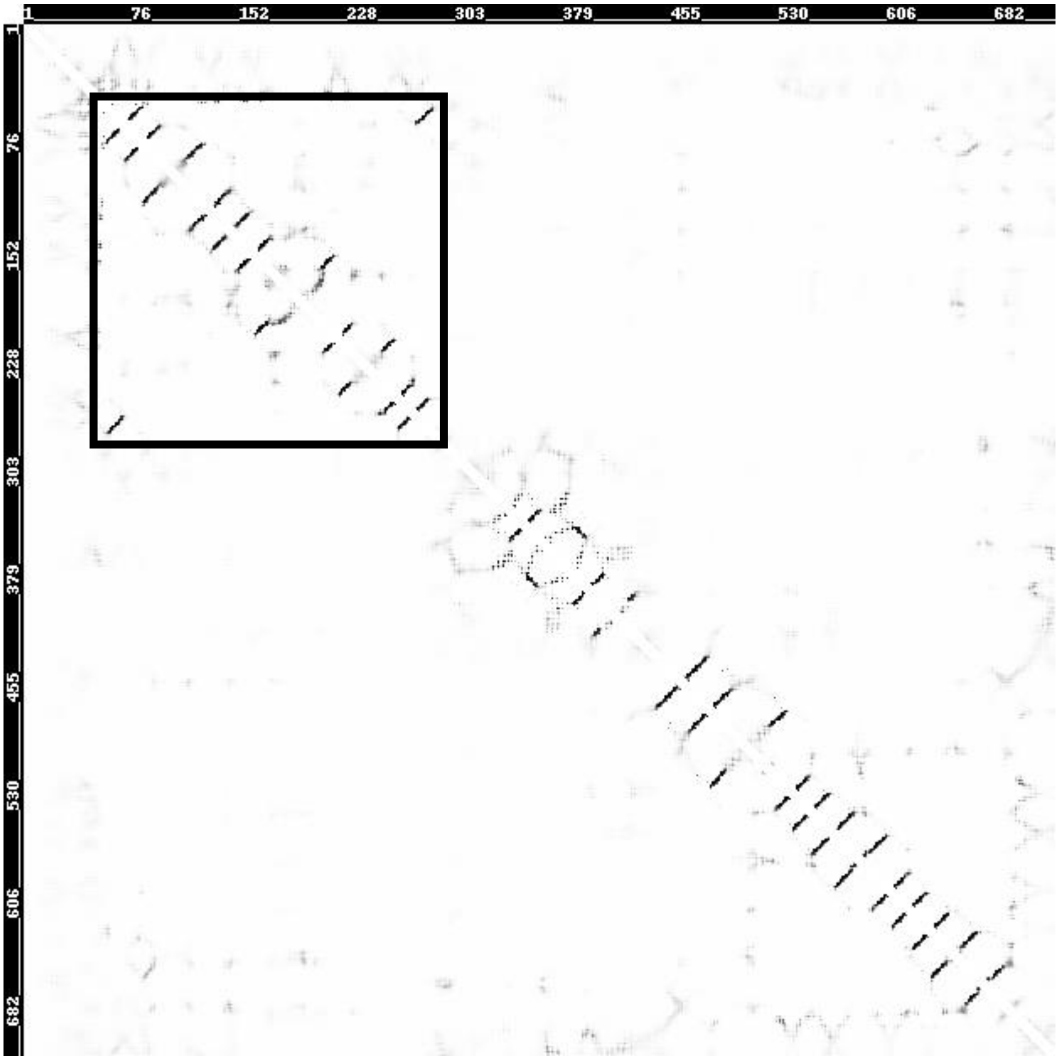


Figure S9. Example of a problematic contact map: family 000, with 4,187 homologous proteins, predicted to form 12-12 architecture. The footprints of a 12-stranded barrel, including the barrel closing signal are clearly available in the N-terminal. However, the signals for a C-terminal barrel domain are less clear. There are 12 signals of contact between adjacent strands, pointing to 13 strands, and no closing signal.

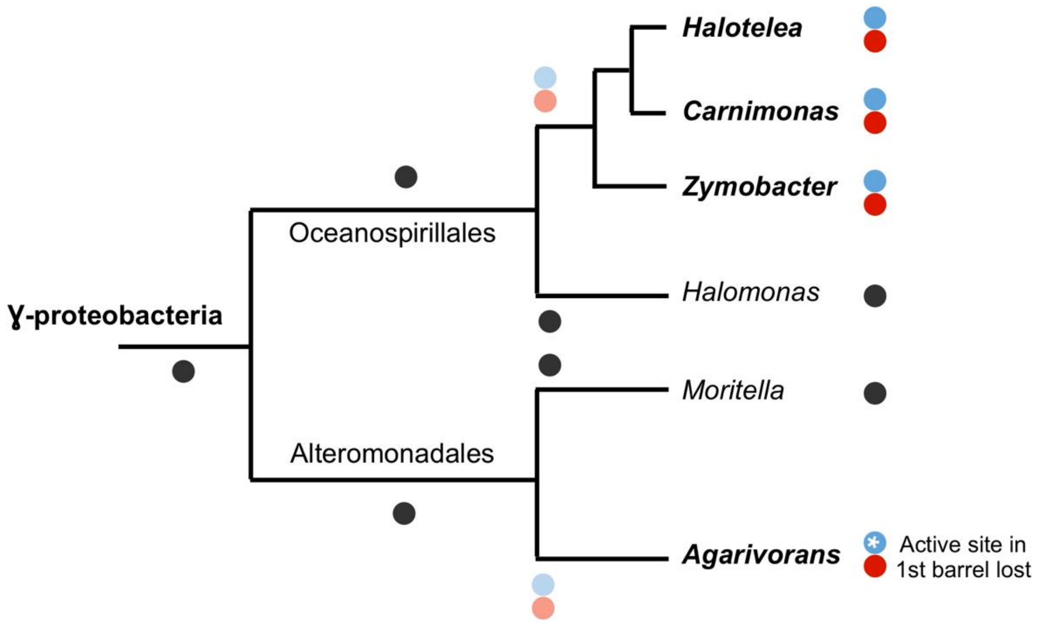


Figure S10. An evolutionary scenario for the independent, parallel emergence of PLA1-PLA1 proteins in the two *Ocenospirillales* and *Agarivorans* lineages. The dots mark the barrels domains – a single dot marks a protein with a single beta barrel, and two dots mark a double-barrel protein, with blue for the N-terminal barrel and red for the C-terminal barrel.

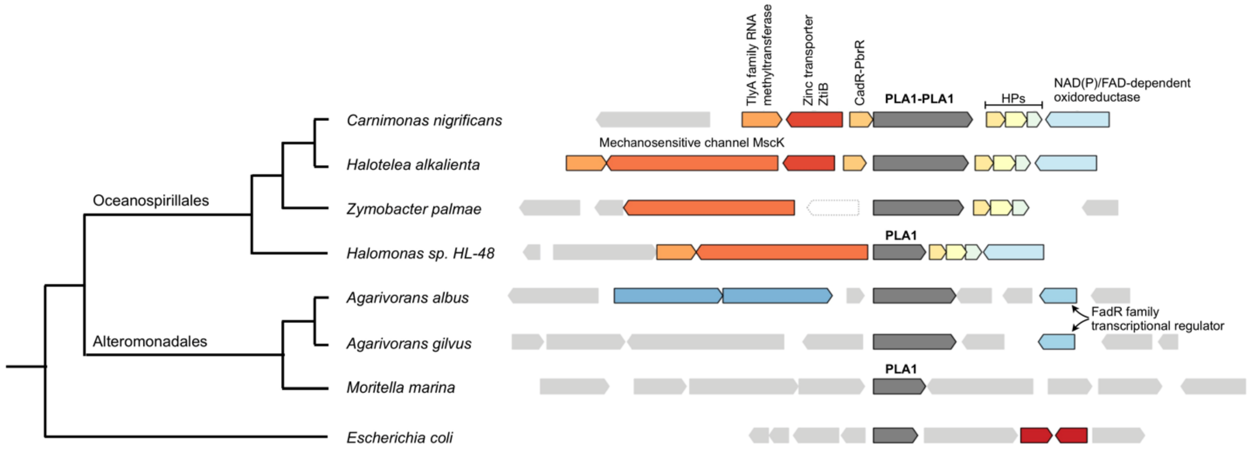


Figure S11. The genomic context of PLA1-PLA1 proteins in comparison to that of their close homologs in two related species (Halomonas and, Mortella marina) and *E. coli.* Homologous genes are colored the same, light grey boxes represent the non-conserved genes, and white boxes the pseudogenes. HP stands for 'hypothetical protein'.

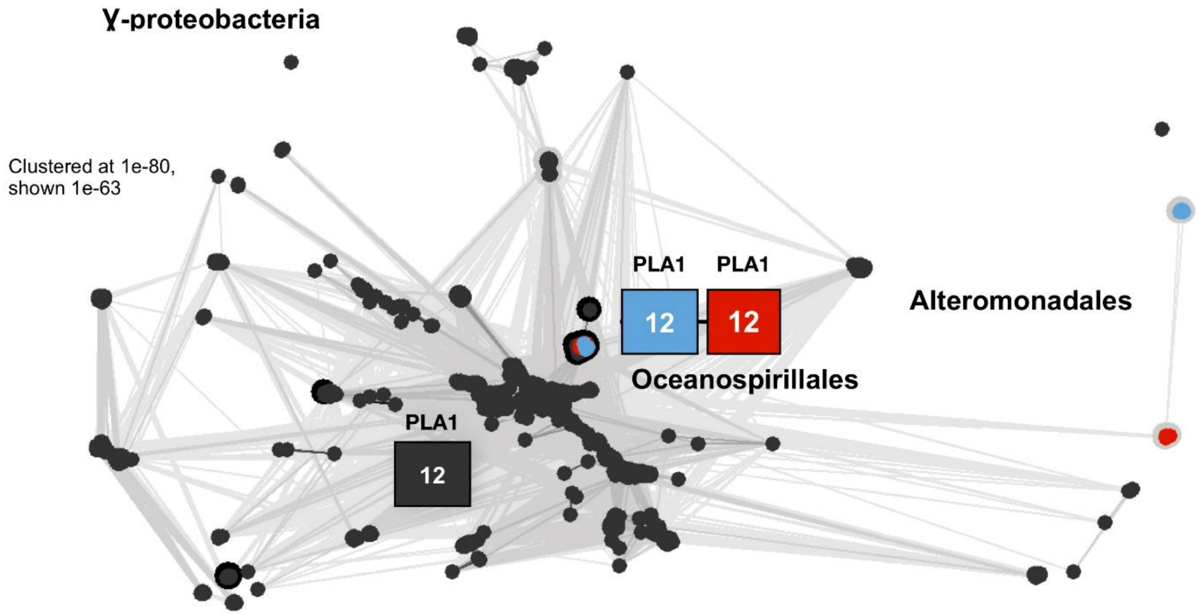


Figure S12. Classification of the PLA1 domains in the six PLA1-PLA1 proteins with their close single barrel homologs in γ-proteobacteria. The dots represent PLA1 barrel domains, and the boxes show the architecture. Black dots represent barrel domains from single-barrel proteins with an architecture of 12, shown in the black box. The blue dots represent the N-terminal barrel domains from the PLA1-PLA1 proteins, and the red dots represent the C-terminal barrel domains from the PLA1-PLA1 proteins. The connected blue and red boxes in the center of the figure represent the 12-12 architecture, whose barrel domains are the red and blue dots near it. The two barrel domains are very similar to each other, and are clustered very close to each other. The barrel domains in *Alteromonadales* are less similar to each other, and are far from each other. Edges connect sequences whose similarity has an E-value of at least $1\times10^{-63}$. Clustering was carried out with CLANS [1] at a p-value of $1\times10^{-80}$ and is shown at $1\times10^{-63}$.

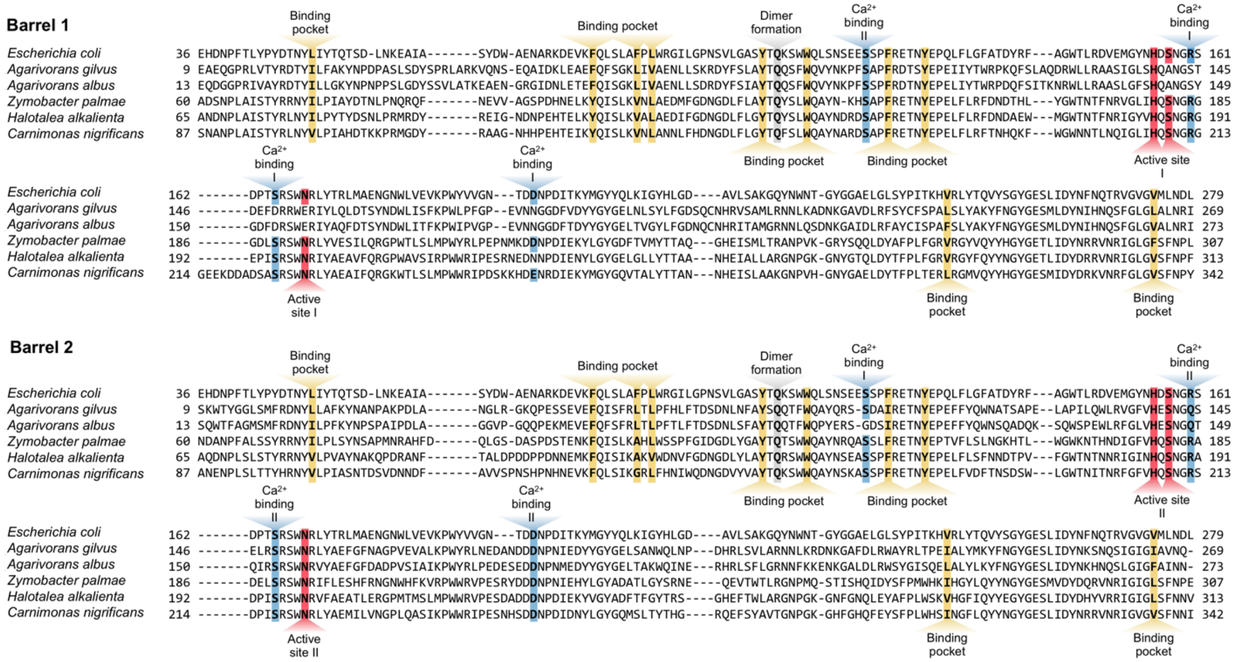


Figure S13. Sequence alignment of the 1st and 2nd PLA1 barrel domains in PLA1-PLA1 proteins to the *E. coli* homolog. Sequence alignment was carried out with PROMALS3D [10], and functionally relevant positions highlighted based on [11].

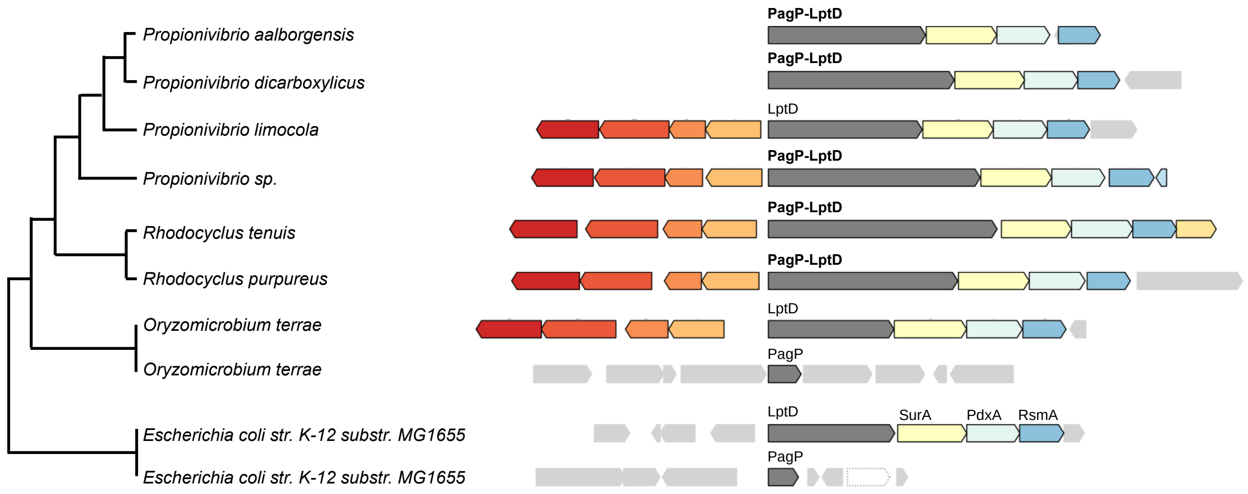


Figure S14. The genomic context of the PagP-LptD MB-family, in comparison to that of the single barrels in *Escherichia coli* (*E. coli*) *K-12* and close species of Propionivibrio. Paralogous genes are colored with the same color, light grey boxes represent the non-conserved genes. Arrow lengths are proportional to the number of the amino acids in the domains.

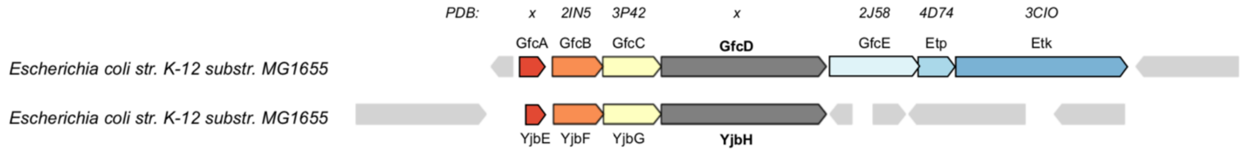


Figure S15. The *gfcABCDE-etp-etk* and *yjbEFGH* operons in *Escherichia coli* (*E. coli*) *K-12*. Homologous genes are colored with the same color and those in the neighborhoods that are not part of the operons are colored grey. The PDB codes for those products for which a full-length or partial structure is known for at least one homolog are shown above the corresponding gene.

| Architecture | #Proteins | PDB Homologs |
|---|---|---|
| 12-12 | 4187 | 5O65_B-4Y25_A fragment |
| 8-8 | 1959 | 2K0L_A-2K0L_A |
| large_single_barrel | 1194 | 6H3I_A fragment-6R2Q_B |
| large_single_barrel | 849 | 6H3I_A fragment-6R2Q_B |
| 8-8 | 813 | 3NB3_A-3QRA_A |
| 8-16 | 469 | 2MLH_A-4GEY_A |
| 8-8 | 392 | 2MLH_A-2MLH_A |
| 8-8-8 | 291 | 3NB3_A-3QRA_A-3QRA_A |
| 22-16 | 273 | 3FHH_A-4AFK fragment-6EHB_B fragment |
| large_single_barrel | 273 | 6H3I_A fragment-6R2Q_B |
| 12-12 | 222 | 5O65_B-very weak 3FIP_B |
| 8-10 | 154 | 2MLH_A-1I78_B |
| 12-12 | 121 | 5O65_B-5O65_B |
| 12-12 | 102 | 6R2Q_B partial-5O65_B |
| large_single_barrel | 100 | 6H3I_A fragment-6R2Q_B |
| 22-8 | 99 | 3CSL_A-2MLH_A |
| 8-8-helical-8 | 91 | 2K0L_A-4RLC_A-5IJN_Z-2X27_X |
| 8-8-8 | 76 | 3NB3_A-6QAM_A-6QAM_A |
| 8-8-8-8-8 | 67 | 3NB3_A-6QAM_A-2K0L_A-6QAM_A-2K0L_A |
| 16-16 | 55 | 4QL0_A-5MDR_F |
| 8-8 | 55 | 2K0L_A-2ERV_A |
| 18-8-8 | 50 | 5ONU_B-2MLH_A-2MLH_A |
| large_single_barrel | 50 | 6H3I_A fragment-6R2Q_B |
| 8-8-8-8 | 45 | 3NB3_A-6QAM_A-2K0L_A-6QAM_A |
| 8-8 | 39 | 3NB3_A-4RLC_A |
| 8-8-helical | 24 | 3NB3_A-6QAM_A-5IJN_Y |
| 12-8 | 23 | 3AEH_B-2MLH_A |
| 8-8 | 23 | 3NB3_A-6QAM_A |
| 8-8 | 19 | 3GP6_A-6QAM_A |
| 12-12 | 17 | 6R2Q_B partial-5O65_B |
| 12-22 | 17 | 3QQ2_A-2FFH_A |
| 22-8 | 17 | 4RDR_A-2MLH_A |
| 12-12 | 15 | 4FQE_A-4FQE_A |
| 8-8-10-8 | 15 | 2MLH_A-2MLH_A-1I78_B-2MLH_A |

| | | |
|---|---|---|
| 8-8-8-8 | 14 | 3NB3_A-2K0L_A-2K0L_A-1QJ8_A |
| 8-8-8-8 | 13 | 2MLH_A-2MLH_A-2MLH_A-2MLH_A |
| 14-8 | 12 | 3PGU_A-2K0L_A |
| 18-8 | 11 | 4AFK_A-2MLH_A |
| 8-8 | 11 | 2MLH_A-2F1V_D |
| 8-8 | 11 | 3NB3_A-2K0L_A |
| 8-10-8 | 10 | 2MLH_A-1I78_B-2MLH_A |
| 8-8 | 10 | 3NB3_A-3NB3_A |
| 12-14 | 9 | 1UYN_X-2X9K_A |
| 12-8 | 8 | 1UYN_X-4FUV_A |
| 8-8-8 | 8 | 3NB3_A-2K0L_A-2MLH_A |
| 10-8 | 7 | 2X55_A-2MLH_A |
| 10-8 | 7 | 1I78-B-2MLH_A |
| 12-12 | 7 | 5O65_B-very weak 6GIE_A |
| 12-8 | 7 | 3QQ2_A-2MLH_A |
| 8-8 | 7 | 2K0L_A-1QJ8_A |
| 8-8 | 7 | 2K0L_A-3QRA_A |
| 8-8-helical | 7 | 3NB3_A-6QAM_A-4PX7_A |
| 12-12 | 6 | 1QD5_A-1QD5_A |
| 8-8 | 6 | 2K0L_A-6QAM_A |
| 8-8 | 6 | 2MLH_A-2MLH_A |
| 8-8 | 6 | 1QJ8_A fragment-2MLH_A-2MLH_A |
| 8-8-8-8-8-8 | 6 | 3NB3_A-2K0L_A-6QAM_A-3QRA_A-2K0L_A-2K0L_A |
| 10-8 | 5 | 1I78-B-2MLH_A |
| 12-12 | 5 | 1TLY_A-1TLY_A |
| 12-12 | 5 | 5O65_B-4Y25_A fragment |
| 12-22 | 5 | 3AEH_B-2QLB_A |
| 22-8 | 5 | 6HCP_B-2MLH_A |
| 8-10 | 5 | 2MLH_A-2X55_A |
| 8-10-8-8 | 5 | 2K0L_A-1I78_B-2MLH_A-2MLH_A |
| 8-12 | 5 | 3NB3_A-5O65_B |
| 8-8 | 5 | 2ERV_A-2K0L_A |
| 8-8 | 5 | 2MLH_A-2MLH_A |
| 8-8 | 5 | 2MLH_A-2MLH_A |
| 8-8-8 | 5 | 2K0L_A-3QRA_A-2MLH_A |
| 22-10 | 4 | 4AFK_A-1I78_B |
| 22-8 | 4 | 6HCP_B-2MLH_A |
| 22-8 | 4 | 6I97_A-2MLH_A |
| 8-8 | 4 | 3NB3_A-6QAM_A |
| 8-8 | 4 | 2MLH_A-2MLH_A |
| 8-8 | 4 | 2F1V_D-2F1V_D |
| 8-8 | 4 | 3NB3_A-1QJ8_A |
| 8-8 | 4 | 2K0L_A-2X27_X |
| 8-8-8-8 | 4 | 3NB3_A-6QAM_A-1QJP_A-1P4T_A |
| large_single_barrel | 4 | |

Table S1. The data used for Figure S2, i.e., the number of proteins in each MB-family with at least 4 similar proteins. **Left column**: barrel architecture. **Central column**: number of similar proteins. **Right column**: The respective PDB homologs. Matches to only part of the target protein are labelled as “fragment”.
